# Drought duration does not impact soil microbiome resilience

**DOI:** 10.64898/2026.07.22.740089

**Authors:** Sreejata Bandopadhyay, Kaizad F. Patel, Sarah J. Fansler, Sophia A. McKever, Ben Bond-Lamberty, Jianqiu Zheng, Vanessa L. Bailey

**Author notes:** Corresponding author: Sreejata Bandopadhyay; Biological Sciences Division, Pacific Northwest National Laboratory, Richland, WA 99352, USA.

## Abstract

Increasing global droughts exert large but poorly understood effects on the microbial communities and ecology of soil. Microbial communities generally show resilience and return to pre-drought conditions when short-term droughted soils are rewet; soils exposed to long-term drought, however, often show a lag upon rewetting, after which microbial communities may or may not return to their pre-stressed conditions. Though short-term droughts have been widely studied, long-term drought manipulation experiments remain rare, especially those that compare microbial response to short-term and long-term drought in tandem. We conducted a 1000-day drought simulation in controlled laboratory conditions with soil cores collected from a tidal freshwater ecosystem in Washington state, USA, and subsequently exposed them to rewetting for two weeks. We also included short-term (30-day and 90-day) drought and rewet treatments to directly compare microbial community and organic matter responses across drought durations. We found distinct microbial taxa belonging to Firmicutes and Actinobacteria enriched after the 1000-day drought, but not after the short-term droughts. While we hypothesized that the microbial community would recover from a short-term drought after rewetting to resemble pre-drought conditions, our results revealed community dissimilarities between rewet and pre-drought conditions across all drought durations. These findings suggest unique microbial life history strategies within certain microbial phyla that make them successful colonizers during an extended drought period, and the influence of environmental and physiological context on microbial responses to rewetting.

**Importance:** Droughts are increasing in frequency and intensity globally with severe implications for ecosystem services and soil functions. It is important to understand how long-term drought impacts soil microbial communities and organic matter chemistry to better predict future ecosystem responses to sustained moisture deficit conditions. We subjected soils to short-term (30 and 90 days) and long-term (1000 days) drought treatments and subsequently rewetted them to understand microbiome recovery to pre-drought conditions. Our results showed that prolonged drought drastically changes the microbial community and soil organic matter profile compared to short-term drought. While we expected the soil microbiome to recover upon rewetting after short-term drought, our results showed an altered microbiome composition, compared to pre-drought conditions, for both short-and long-term drought, suggesting microbial responses to soil rewetting was independent of drought duration imposed. These results provide important insights into soil biological and chemical functions that remain sensitive to change under fluctuating soil moisture conditions and future drought scenarios.

## Introduction

Droughts are intensifying and increasing in frequency globally [1, 2] with major but uncertain implications for soil ecosystem functions. Most documented droughts in natural environments are short-term and moderate in intensity [3, 4]. However, drought length varies among regions, with the longest lasting droughts reported in Oceania (12 months) and shortest (5 months) in northern Europe [5].

Experimental simulations of drought have largely been short-term [6–10]. Short-term drought causes temporary reductions in soil microbial activity due to water limitation [11], while long-term drought can cause persistent reduction in metabolic processes and microbial diversity [12]. Microbes may enter a dormant state [13–15] slowing down key processes like nutrient cycling and soil organic matter (SOM) decomposition [16]. Modest shifts in microbial community composition are common during short-term drought, resulting in a transient dominance of drought-tolerant microbes [17]. Adaptive mechanisms such as production of osmolytes and extracellular polysaccharides are often used to combat desiccation [17, 18], with long-term drought shown to favor oligotrophic microbes [19, 20]. In extreme cases, sensitive microbes may be completely eliminated, leading to lasting changes in microbial community dynamics [21]. However, the mechanisms linking drought to microbial function remain poorly understood, particularly across different drought durations.

Rewetting the soil, post drought, triggers rapid and often dramatic microbial responses, leading to nutrient pulses and CO₂ bursts (the “Birch Effect”) [22, 23]. For the short-term, microbial functionality commonly recovers with rewetting [24]. Copiotrophic bacteria such as Proteobacteria and Bacteroidetes generally recover quickly after rewetting [25] as water availability enhances microbial respiration and nutrient uptake. However, the recovery trajectory varies, such that copiotrophs can decrease after drought and rewet in a pH dependent manner [26]. In contrast, soils exposed to long-term press disturbance or intense drought often show incomplete recovery [27], and depend on legacy [28], with lag times to microbial growth and respiration after rewetting [29], indicating reduced resilience of the system.

Critically, the impact of drought duration on the ability of the microbial community to recover, and whether there is an inflection point beyond which communities are unable to recover to pre-drought state, remains unclear. While short-and long-term responses to drought and rewet are reported as separate studies in the literature, a comparison of these responses using concurrent short-and long-term drought-rewetting events in the same study system remains underexplored, especially those that deploy droughts for more than two years. Such prolonged drought studies are rare in the literature, making it difficult to identify comparative, long-term response patterns.

To improve our understanding of microbiome response to long-term drought, we conducted a comparative study of drought-rewetting effects on soil microbiome dynamics over the short-term (30 days, 90 days) and long-term (1000 days). This report constitutes one of very few controlled laboratory experiments, if any, that simulated drought conditions for ∼3 years and used parallel short-term drought-rewetting treatments as part of the same study system. We hypothesized that 1. Soil microbial community shifts over 1000 days will select taxa not prevalent during short-term drought. 2. Soils that are dried longer will experience winnowing of the revivable microbial population and thus will show less differentiation between drought and drought+rewet samples (henceforth, d+rewet). 3. Soils exposed to short-term drought (Day 30, Day 90) would become more similar to pre-drought (Day 0) upon rewetting, compared to long-term drought (Day 1000).

## Methods

### Site description and sampling

Soil samples were collected from the poorly drained floodplains along the Secret River (46.308° N, 123.690° W), a tributary to Grays Bay in Washington, USA [18, 30]. This tidal freshwater system features an extensive network of tidally filled underground channels that fill and recede daily. Vegetation at the site is dominated by Sitka spruce (*Picea sitchensis*) and the soils are classified as Ocosta silty clay loam (fine, mixed, superactive, acid, isomesic Sulfic Endoaquepts). Soil characteristics are reported in Table S1.

Intact soil cores (5 cm diameter, 15 cm height) were sampled in April 2019, placed in a cooler on blue ice, and brought immediately to the Pacific Northwest National Laboratory in Richland, WA, USA, arriving within 24 hours of sampling. The cores were stored at 4°C until ready for incubation.

### Drought incubation

The cores were dried one of two ways: (a) air drying – the cores were air-dried at room temperature until they reached constant weight, representing a passive/less intense drying; and (b) forced drying , to simulate more intense and accelerated drying conditions – the cores were dried using diurnal cycles of 12 hours at 35°C and bright light (987 µmol light), followed by 12 hours at 21°C in the dark, until the soils achieved constant weight.

The dried cores were then held at room temperature for 30, 90, or 1000 days to simulate extended drought of varied durations. At the end of each drought period, the cores were either (a) deconstructed and extracted immediately (drought treatment), or (b) allowed to saturate for two weeks (d+rewet treatment, wetting from below in a water bath) and then deconstructed and processed for extractions. An additional set of cores (time-zero) was deconstructed and processed for extractions at the start of the experiment, using field-moist cores. Individual time points of sample collection henceforth referred to as Day 0, Day 30, Day 90, and Day 1000.

### Soil core deconstruction

During deconstruction, the cores were split into the top 5 cm and remainder 5cm-end portions, and the two depths were processed separately. All samples were sieved through 4 mm mesh and homogenized. The material was subsampled for microbiome analysis (16S rRNA gene sequencing) and organic matter characterization (non-targeted FT-ICR-MS and ^1^H-NMR).

### Soil DNA extractions and quantification

DNA extractions were performed using the Quick-DNA Fecal/Soil Microbe Kits (Zymo Research, Irvine, CA, USA) as per manufacturer’s instructions, with one modification — 0.25 g of wet soil sample was mixed with a BashingBead^TM^ buffer and bead beating was done for ∼20 minutes on a vortex-genie to ensure that tough-to-lyse bacteria and their spores were accurately captured in the sequencing results. This was particularly critical for the Day 1000 drought samples. DNA was quantified using the Qubit dsDNA BR assay kit on a Qubit Flex fluorometer (Invitrogen, Carlsbad, CA, USA).

### 16S rRNA (V4) gene amplicon sequencing

Amplicon sequencing was completed targeting the V4 region of the 16SrRNA gene using 515F (forward primer with index barcode) and 806R primers [31–33]. Amplicon concentrations were determined using Quant-iT PicoGreen dsDNA assay kit (Invitrogen, Waltham, MA) following the manufacturer’s recommendation. Sequencing was completed on a Illumina MiSeq platform; a v2 500 cycle kit was used with a 15% PhiX spike-in. The MiSeq was used to generate fastq files by demultiplexing the data as a final sequencing output. Further details on the sequencing protocol, including PCR reactions set up and thermocycler parameters can be found in the Supplementary Information.

### 16SrRNA gene sequence analysis

All sequences were processed as described in [8]. Briefly, sequences were analyzed using QIIME2 with all paired-end sequences compressed and denoised using the DADA2 plugin. The denoising step ensured dereplication of sequences, filtering of chimeras and merging of paired-end reads. FIGARO was used to determine the truncation parameters which were used as input within the DADA2 plugin. The truncation length was set to 102 F and 183 R for all data with minimum overlap set to 30 base pairs which resulted in 93% merging success. All truncation was performed from the 3’ end to ensure consistent final read lengths. Representative sequences were classified using SILVA v138 to generate the taxonomy file. The resulting ASV and taxonomy files were exported to R for ecological analysis.

### Microbiome data analysis and ecological statistics

All microbial data analyses and visualization were performed in R version 4.5.1 [34]. Contaminants (using R package *decontam* [35]), mitochondria, and chloroplast were removed prior to downstream processing of ASV tables. Diagnostic results from contaminant analysis along with rarefaction curves are reported in Figure S1. Beta diversity was calculated using Bray-Curtis distances and visualized using a principal coordinate analysis (PCoA). Permutational analysis of variance (PERMANOVA) was computed using the adonis function in *vegan* [36] package to analyze significant differences in Bray-Curtis distances between experimental variables (factor: saturation, levels: timezero, drought, drought+rewet; factor: time, levels: Day 0, Day 30, Day 90, Day 1000; factor: depth, levels: 0-5cm, 5cm-end). Alpha diversity was evaluated using the estimate_richness function in R package *phyloseq*. Both the number of observed ASVs and the Inverse Simpson diversity index were computed and visualized.

Indicator species analysis was performed using the *indicspecies* package [37] in R for each drought time point. We used the abundance-based counterpart of Pearson’s phi coefficient of association within the multipatt function. To correct the phi coefficient for unequal group sizes the “func” parameter within multipatt was set to “r.g.” P values were adjusted for false discovery rates. Overlapping and shared indicators across drought time points were visualized using a venn diagram using the venn.diagram function from the *VennDiagram* package in R [38].

Phylogenetic trees were plotted using a rooted tree generated from the representative sequences in QIIME2 using the align-to-tree-mafft-fasttree pipeline. Bray-Curtis distance heatmaps were used to visualize the dissimilarity index between samples (drought vs d+rewet) and their temporal patterns across Day 30, Day 90 and Day 1000.

Relative abundance of indicator taxa was also plotted using heatmaps. All heatmaps were generated using the pheatmap function within the *pheatmap* package in R [39]. ANOVA and Sidak HSD posthoc tests were computed to assess differences in alpha diversity metrics (richness and diversity). Differential expression analysis was performed on unrarefied read count data using *DESeq2* package [40] to cross check taxa that were differentially abundant at Day 1000 versus Day 30 and Day 90, to confirm that rarefaction did not impose a bias towards the taxonomic classes identified using indicator species analysis.

### Water extractable organic carbon (WEOC) characterization, data analysis, and statistics

We used Nuclear Magnetic Resonance (NMR) and Fourier-transform ion cyclotron resonance mass spectrometry (FT-ICR-MS) to characterize WEOC. Experimental details, with further processing using the *nmrrr* [41] and *fticrrr* [42] workflows in R, are provided in the Supplementary Information. We used permutational multivariate analysis of variance (PERMANOVA) and principal components analysis (PCA) to analyze FT-ICR-MS and NMR relative abundance data. Statistical significance was determined at α = 0.05 using ANOVA and PERMANOVA. Because there was no significant effect of drying type (air drying vs. forced drying) on the WEOC chemistry, we focus primarily on the air-dried samples. All biochemical data analyses and visualization were performed in R version 4.5.1 [34], using primarily the *dplyr* v1.0.1 [43] and *vegan* v2.5-6 [44] packages for data processing and analysis, and *ggplot2* v3.3.2 [45] and *PNWColors* [46] packages for data visualization.

## Results

### Microbial communities differed by saturation and time with the highest richness observed at Day 1000 drought

#### Beta diversity

Bray-Curtis distances between Washington soil samples revealed significant differences by saturation type (PERMANOVA Pseudo-F=13.98, p<0.001, R^2^=0.19), time (PERMANOVA Pseudo-F=7.61, p<0.001, R^2^=0.10) and depth (PERMANOVA Pseudo-F=3.38, p<0.001, R^2^=0.02) (Figure 2A, Table S2). Significant interaction effects were observed between saturation and time (Pseudo-F=5.47, p<0.001, R^2^=0.08), saturation and depth (Pseuodo-F=1.51, p<0.05, R^2^=0.02) and time and depth (Pseudo-F=1.54, p=0.03, R^2^=0.02). No significant differences in microbial community composition were observed between air dry and forced dry methods and hence drying was not tested as a separate factor in subsequent analyses.

**Figure 1:**
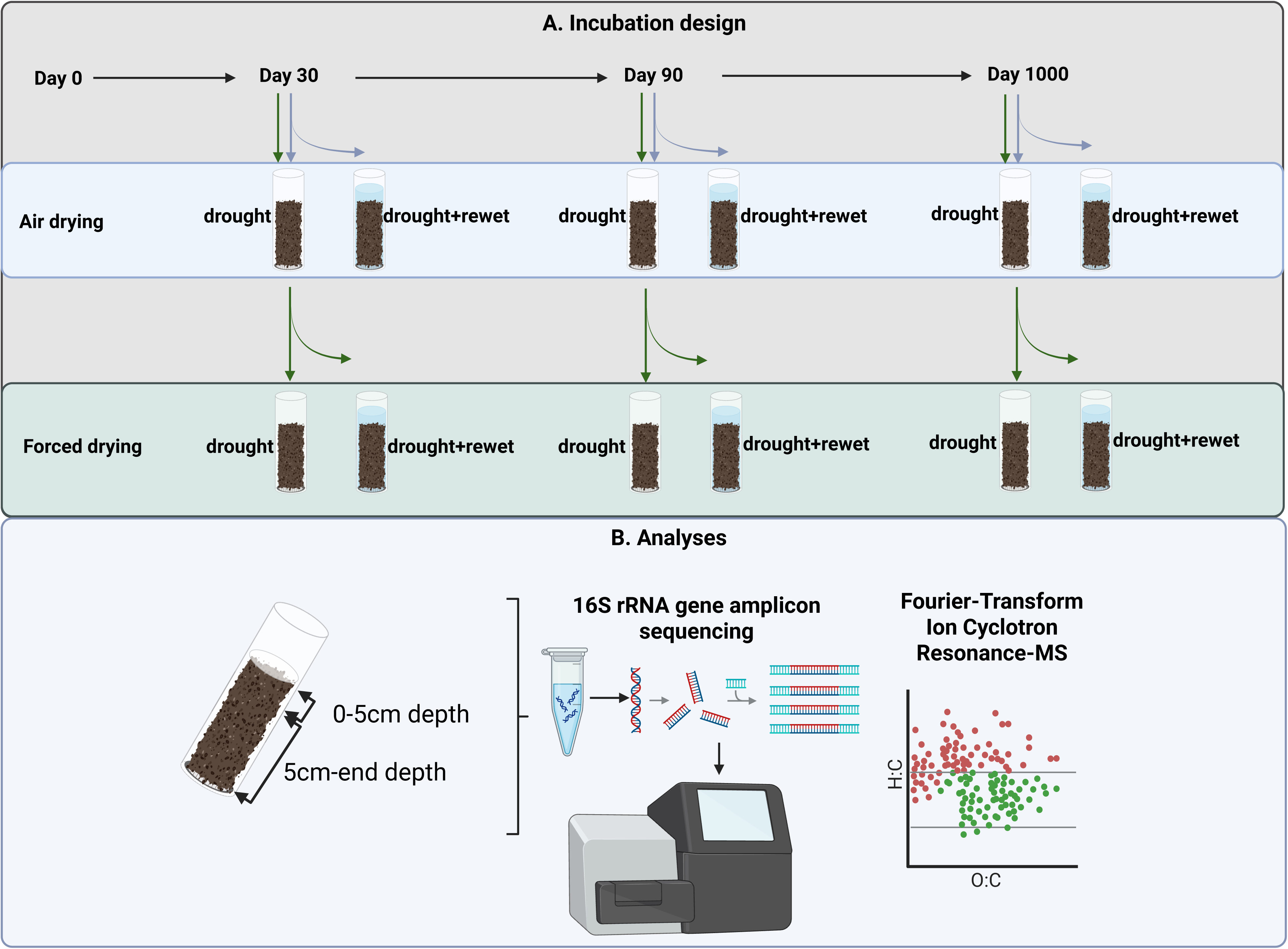
Experimental design and set up. Created in BioRender. Bandopadhyay, S. (2026) https://BioRender.com/3ypl0oe **ALT TEXT:** Graphical representation of the experimental design and set up used in the study.

**Figure 2:**
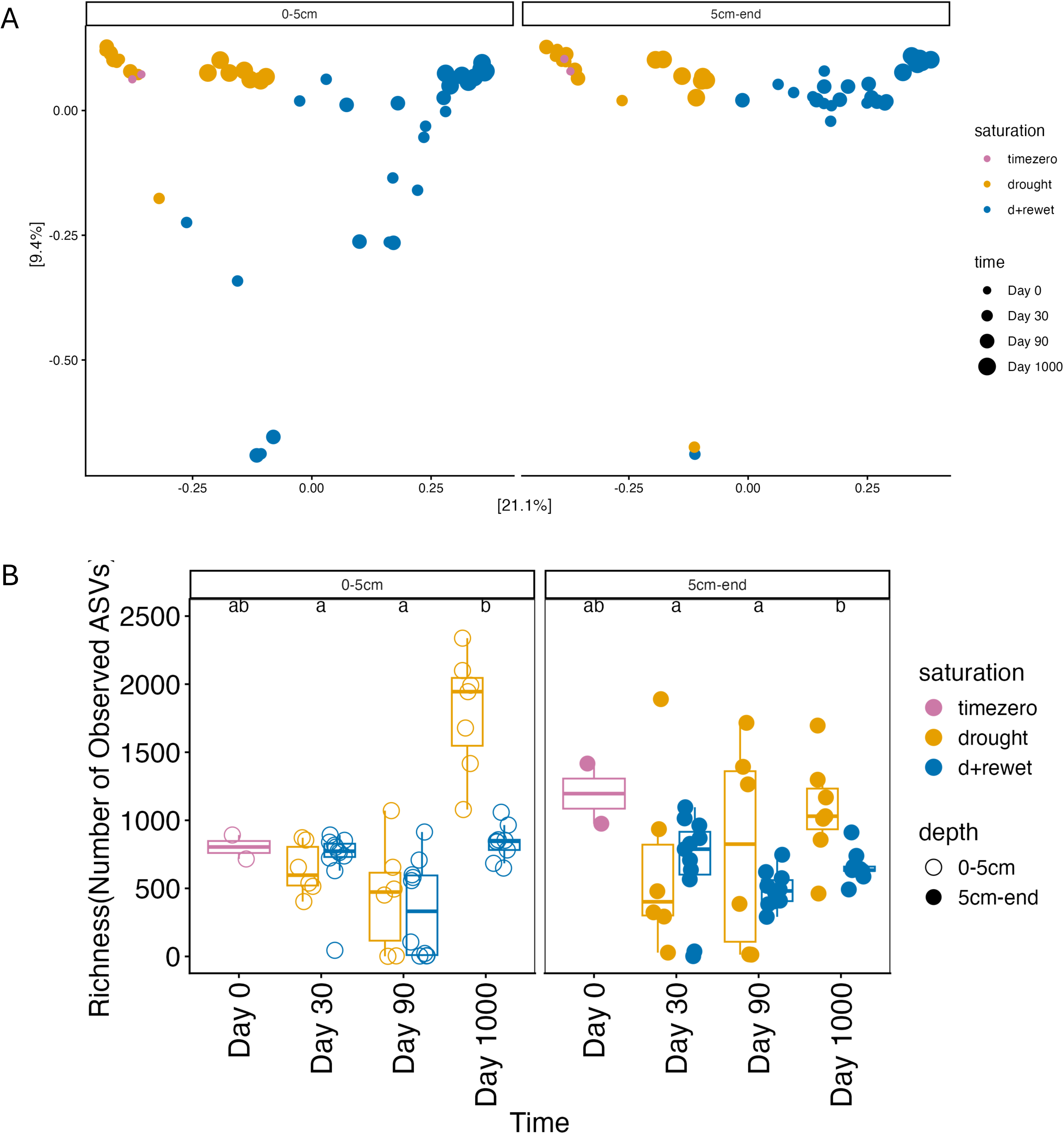
Microbial communities differed by saturation and time, with highest richness observed at Day 1000 drought. A. Differences in microbial community composition by treatment (saturation) and time across both 0-5 cm (left panel) and 5cm-end depth (right panel) using a principal coordinate analysis (PCoA). B. Microbial community richness denoted as number of Observed Amplicon Sequence Variants (ASVs) across all time points, treatments and soil depths for Washington soils. The whiskers in the boxplots represent 1.5 × the Interquartile Range (IQR). **ALT TEXT:** Graphical representation of the differences in microbial community composition, and richness, by treatment, time and soil depth across all samples used in the study.

#### Alpha diversity

The highest richness (number of observed Amplicon Sequence Variants (ASVs)) was seen in the Day 1000 drought samples for both 0-5 cm and 5cm-end depths (Figure 2B). For the 5cm-end depth profile, the Day 1000 drought samples reflected similar richness values as the Day 0 samples (Tukey post hoc test p>0.05). The shallower 0-5cm depth had greater richness for the Day 1000 drought samples compared to the Day 0 samples (Tukey post hoc test p=0.002).

### Total microbial community dynamics across drought durations and saturation levels

#### Effect of time on drought and d+rewet samples

Microbial community composition over time showed enrichment of endospore-forming members within the phylum Firmicutes, specifically Clostridia and Bacilli at Day 1000 compared to Day 30 and Day 90 timepoints (Figure 3) in Washington soils for both drought and d+rewet samples (Dunn’s test p<0.05). Exospore-forming Actinobacteria were enriched at Day 1000 drought versus Day 90 (Dunn’s test p<0.001) and Day 1000 drought versus Day 30 (Dunn’s test p=0.002) but there were no differences in Actinobacteria abundance across time points in d+rewet samples (Dunn’s test p>0.05). Alphaproteobacteria decreased in relative abundance at Day 1000 time point compared to Day 30 (Dunn’s test p<0.05) and Day 90 samples (Dunn’s test p<0.05) for drought and d+rewet treatments. Gammaproteobacteria also decreased in relative abundance at Day 1000 compared to Day 30 and Day 90 (Dunn’s test p<0.05) but only for the drought treatment. Effect of time across the whole community is reported in Table S2.

**Figure 3:**
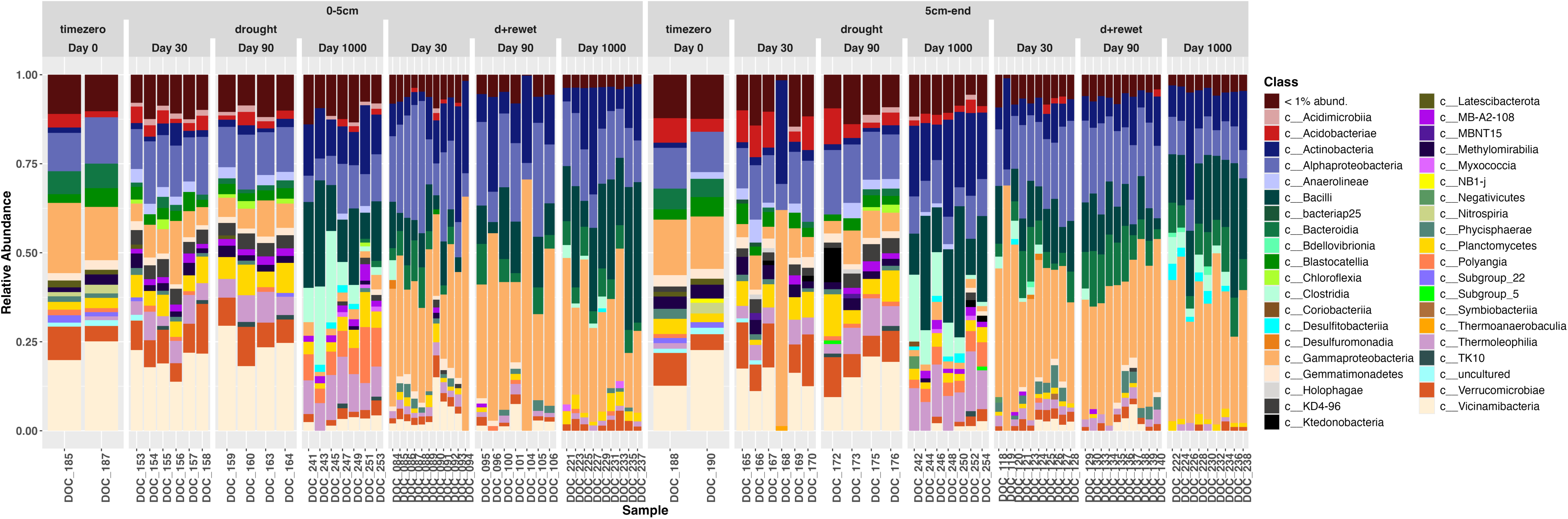
Microbial community composition across all time points and treatments. Stacked barplot shows relative abundances at the Class level. Taxa whose abundances were less than 1% are aggregated to a single color (deep brown) at the top of the barplot. **ALT TEXT:** Graphical representation showing the different microbial classes and their relative abundances for each sample across all treatments, time points, and soil depths used in the study.

#### Effect of Saturation

Gammaprotebacteria increased in d+rewet soils compared to drought soils (Kruskal Wallis rank sum test p<0.001) across all time points whereas Thermoleophilia decreased in d+rewet samples compared to drought samples across all time points (Figure 3, Kruskal Wallis rank sum test p<0.001). Acidobacteriae decreased in abundance or were lost across all time points in the d+rewet samples compared to the drought samples (Kruskal Wallis rank sum test p<0.001).

Within the Firmicutes phylum, Bacilli did not differ in abundance between drought and d+rewet samples at Day 1000, but abundance of Clostridia was significantly higher in Day 1000 drought compared to d+rewet samples (Kruskal Wallis rank sum test, p=0.002). Similarly, Actinobacteria increased in abundance in the Day 1000 drought samples compared to the d+rewet samples (Kruskal Wallis rank sum test, p=0.03).

However, for the shorter durations, both Day 30 and Day 90, Bacilli, Clostridia and Actinobacteria were all significantly higher in the d+rewet samples compared to the drought samples (Kruskal Wallis rank sum test, p<0.05) except for Clostridia abundances at Day 90 which were not significantly different between drought and d+rewet samples. The effect of saturation across the whole community is reported in Table S2.

### Indicator taxa for drought at Day 1000 show greater persistence after rewetting compared to earlier time points

We conducted an indicator species analysis by drought duration (across Day 0, Day 30, Day 90 and Day 1000) (Figure 4A, Venn diagram). 36 ASVs were unique to Day 0 and were not observed in the drought samples. Two ASVs were shared across all drought samples but not Day 0 – these belonged to phylum Actinobacteria (class MB-A2-108, and Thermoleophilia). Seven indicators were shared between Day 30 and Day 90 drought time points which were lost at Day 1000, and 48 unique taxa were detected after Day 1000 of drought. Overall, the indicator taxa gained at Day 1000 drought showed greater persistence after rewetting compared to indicator taxa for Day 30 and Day 90 drought time points, for which rewetting caused loss of several indicator species (Figure 4B). A list of all shared and unique indicator taxa is provided in Table S3 along with a heatmap showing the relative abundance of all the 48 unique indicator taxa for Day 1000 drought across time points and saturation levels (Figure S2). Drought responsive indicator taxa were not sensitive to rarefaction (Figure S3A, B) and generally showed phylogenetically conserved responses at deeper taxonomic levels (Figure S4A, B).

**Figure 4:**
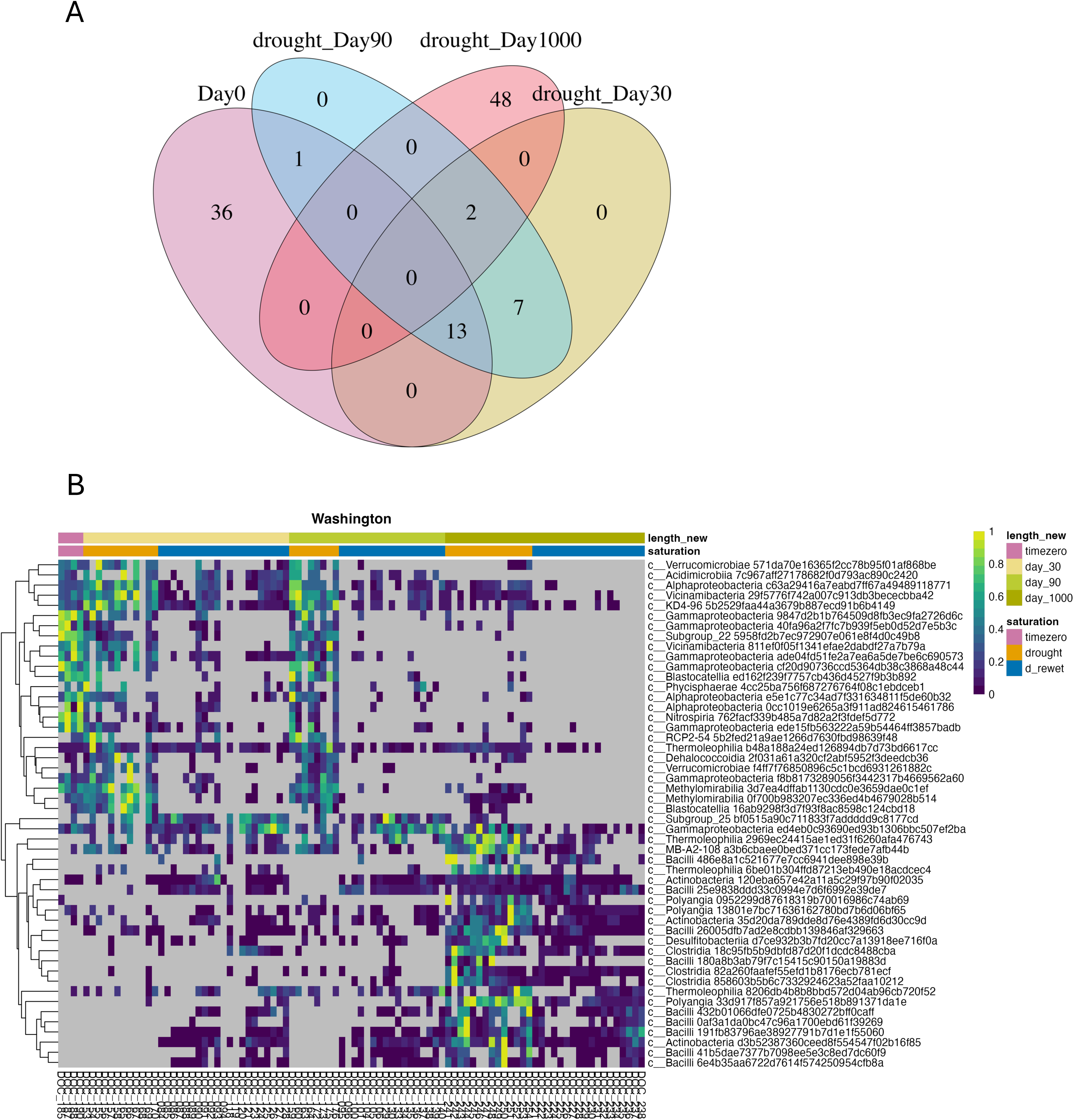
Indicator ASVs unique and shared between short-term and long-term drought. A. Indicator taxa Venn diagram by drought duration. Overlapping numbers show indicator ASVs shared across drought durations, unique ASVs associated with a specific drought duration are indicated in the center of the ellipses. B. Heatmap showing the relative abundances of the top 50 most abundant indicator ASVs across all treatments and timepoints. Abundances are scaled for each ASV using a maximum standardization approach. Color gradient denotes relative abundance values (yellow: high abundance, blue: low abundance, grey: Abundance==0). **ALT TEXT:** Graphical representation of the indicator taxa that are unique and shared across the different drought time points, with the top 50 most abundant indicator taxa picked to represent their relative abundances across time zero, drought and rewet treatments.

### Microbial communities were more similar between drought and d+rewet for long-term drought (Day 1000) compared to short-term drought (Day 30 and Day 90)

We compared the Bray-Curtis dissimilarities between drought and d+rewet communities for each time point (short-and long-term drought) and for each soil depth separately (Table S4, Figure S5). All comparisons between drought and d+rewet samples were significant (PERMANOVA p<0.05 across all drought and d+rewet comparisons), indicating a change in community composition between drought and d+rewet samples for all time points and soil depths. However, differences between drought and d+rewet samples showed greater variability for the earlier time points (Day 30 and Day 90) and were relatively higher (greater dissimilarity between drought and d+rewet) compared to Day 1000. For the Day 1000 time point, the differences between drought, and d+rewet samples were largely consistent across samples, leading to a greater R^2^, P and F value for both depth profiles analyzed (0-5 cm, and 5cm-end, Table S4).

### Soil organic matter composition was distinct at Day 1000 drought and d+rewet conditions compared to Day 30 and Day 90 time points

Soil organic matter showed a distinct profile of organic molecular classes for long-term drought and d+rewet treatments (Figure 5A). Day 1000 (long-term drought) formed a distinct cluster separated from Day 0, Day 30 and Day 90. There was no separation between drought and d+rewet samples for short-term drought though some separation was observed for Day 1000. Aggregating results across the two depths, time of drought duration accounted for 47% of total variability and saturation (drought vs. d+rewet) accounted for 35% of variability (Table S5). Unsaturated/lignin-like molecules were higher at Day 1000 drought and d+rewet compared to short-term drought, an observation consistent across both soil depths analyzed (Figure 5B). Van Krevelen diagram, with molecular classification of the peaks identified via FT-ICR-MS (Figure S6) shows each point representing a unique peak, plotted against its molecular H/C and O/C ratios.

**Figure 5:**
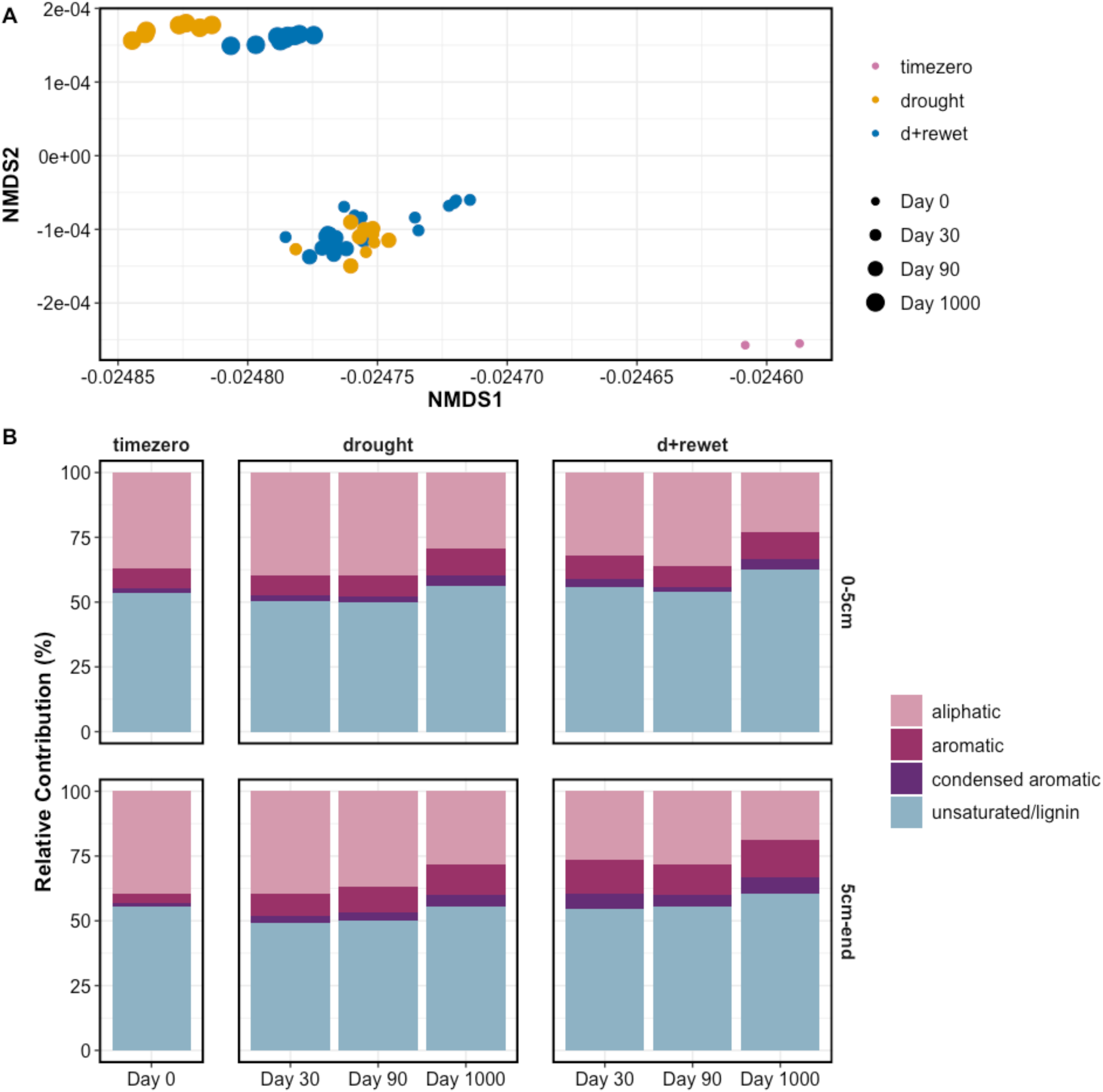
Soil organic matter profiles across short-and long-term drought and d+rewet conditions, and their comparison to Day 0 samples before drought was initiated. A. NMDS biplot of FTICR-MS profiles across all timepoints and treatments. B. Stacked barplot showing relative abundances of organic compounds classes. Relative abundances reflect aggregated responses at the level of individual molecular formulae for each compound class with each molecular feature having a presence/absence value instead of true abundances. **ALT TEXT:** Graphical representation of the differences in soil organic matter composition across treatments and time points, with a visual showcasing the relative abundances of the different organic compound classes by treatment, time point and soil depth.

Saturation (drought vs. d+rewet) accounted for 39% of total variability in the NMR spectra of samples (Table S6). There was a lot of inter-replicate variability, but some general trends emerged. Day 0 soils were greater than 50% aliphatic, and drought samples showed presence of alpha-H groups (Figure S7). Drought and rewet (d+rewet) treatment showed an increase in aromaticity, with loss of alpha-H groups.

### Prolonged drought for 1000 days increased dissimilarity to pre-drought condition compared to short-term drought, though rewetting results in an altered community composition in both cases

We found that the Bray-Curtis dissimilarity to pre-drought states were higher in the d+rewet condition compared to their dissimilarities during drought over both the short-term and long-term drought conditions (Figure 6). During short-term drought (Day 30 and Day 90), communities were more similar to pre-drought conditions with the dissimilarity index increasing during long-term drought (Day 1000) (linear model R^2^=0.68). Drought and rewet (d+rewet) conditions showed more stability in their dissimilarity to pre-drought conditions over the short-and long-term, though with higher dissimilarity index compared to drought condition across all time points (linear model R^2^=0.33). The group means for drought and d+rewet conditions were significantly different (analysis of covariates controlling for time, ANCOVA p<0.001, Figure S8).

**Figure 6:**
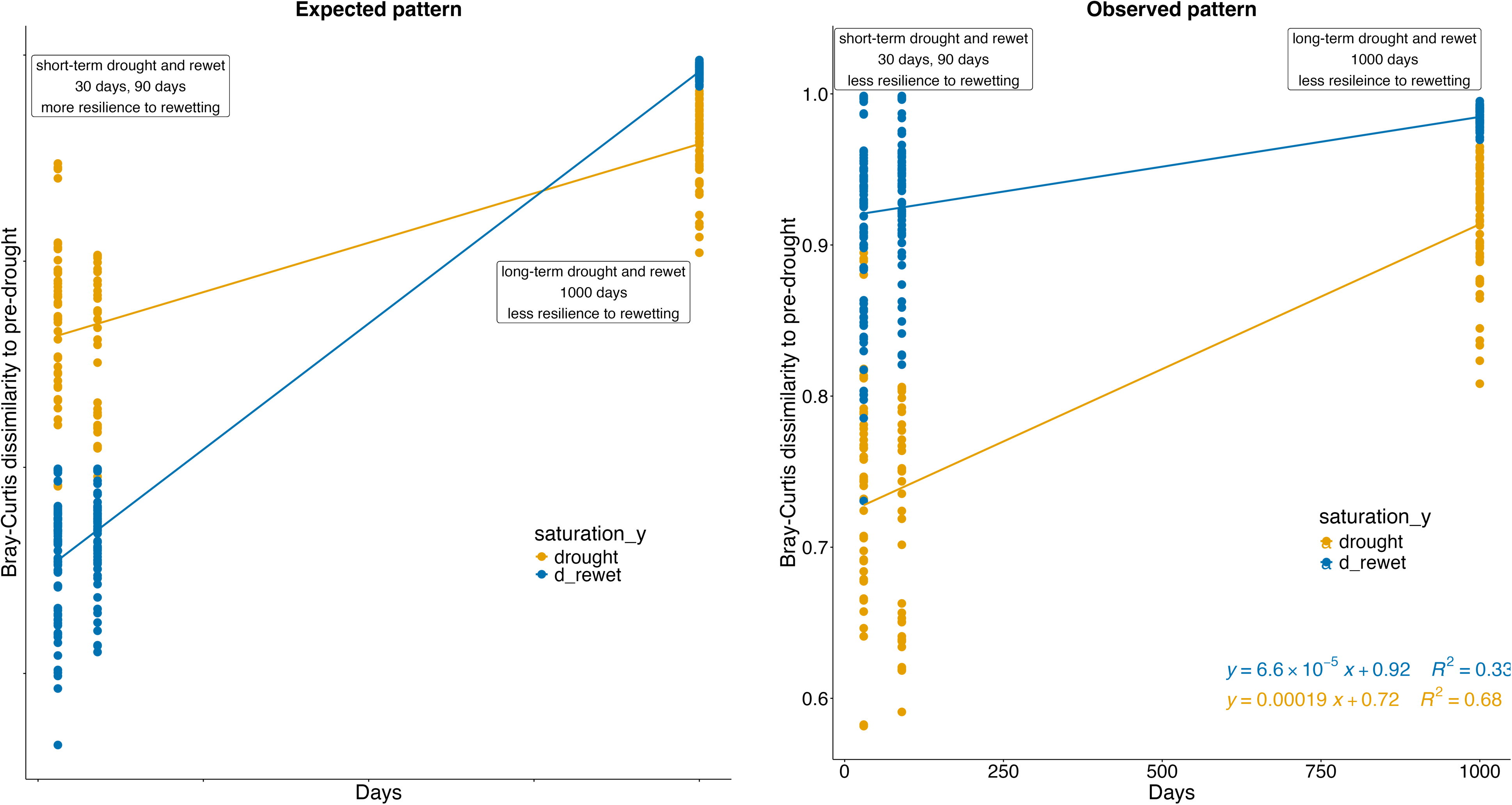
Expected and observed temporal patterns of Bray-Curtis dissimilarity between drought/d+rewet and pre-drought conditions. Expected data includes observed data for the drought treatment and hypothesized (mock) data for the d+rewet treatment as per Hypothesis 3. **ALT TEXT:** Graphical representation of the hypothesized (expected) versus observed patterns in dissimilarity indices between drought/rewet and pre-drought conditions.

## Discussion

### Prolonged drought caused distinct compositional changes in soil microbial communities and organic matter chemistry

We observed an increase in Actinobacteria and decrease in Alphaproteobacteria from Day 30 to Day 1000 drought, consistent with previous work [47–49]. The increased alpha diversity we observed at Day 1000 is supported in the literature [49] where soils with a history of drought have shown higher overall bacterial alpha diversity at the end of a drought treatment. This suggests the importance of soil legacy which may impact microbial adaptation.

Acidobacteria and Verrucomicrobia have been shown to exhibit opportunistic behavior during drought conditions, with a rapid decline in and fast response to ribosome synthesis during drought and subsequent rewet [50]. Our results support these prior findings, with Acidobacteria and Verrucomicrobia decreasing as drought extended to Day 1000 and a continued persistence of Verrucomicrobia across all time points after rewetting (Figure 3). Actinobacteria exhibited an opposite pattern in [50], with ribosome synthesis stimulated during drought and reduced after rewetting, supporting our observation of peak Actinobacteria abundance at Day 1000 drought. Such an accumulation of transcriptional machinery during drought might enable Actinobacteria to invest in growth when conditions become favorable for nutrient acquisition.

Firmicutes, specifically *Bacilli* and *Clostridia* sp., had a similar response as Actinobacteria in our study, increasing in relative abundance with drought duration and reaching peak abundance at Day 1000. Enrichment of Bacilli and decline of Acidobacteria during drought was also reported in [51]. This is contrary to the life strategy proposed in [50] where Firmicutes, Proteobacteria, Chloroflexi, and Planctomycetes showed a more resistant response pattern with a stable ribosome content throughout drought and rewet conditions. The drought duration in [50] was 5 months of dry-down followed by a 2 hr rewet, falling within the short-term drought range for our study. Over the short-term (∼3 months), we observed a relatively stable response of Firmicutes with abundances ranging from very low read counts to no detectable reads. The lack of read counts for Firmicutes during short-term drought should not be misconstrued as an absence of the group but rather an artifact of sequencing depth. Overall, we found phylogenetically conserved bacterial responses to long-term drought at the phylum level which is consistent with previous studies [50, 52–54].

We observed a dramatic shift in soil organic matter chemistry at Day 1000 for both drought and d+rewet as compared to the earlier time points (Figure 5) indicating a change in carbon allocation during water limitation. Previous studies focused on SOM characterization during drought via FTICR-MS have reported greater relative amounts of lignin-like versus condensed aromatic polyphenol formulae, consistent with our results and suggesting reduced decomposition of SOM and microbes under stress [55, 56]. The increase in alpha-H groups in drought samples’ NMR spectra can be indicative of protein content and could be linked to necromass. However, this result was opposed to other studies that demonstrated decrease in labile protein-like compounds during drought due to preferential consumption of bioavailable compounds during stress and decreased amino acids and carbon allocation to biomass [56]. In our study, microbial cell lysis during drought likely released microbial necromass, which was then rapidly consumed when rewet as demonstrated by the loss of alpha-H groups. This observation is supported by a study [57] that observed significantly increased bacterial necromass carbon in topsoil and subsoil layers during moderate and intensive drought treatments from a 3-year field experiment in a forest plantation, though these results were obtained from gas chromatography versus NMR.

### Widespread enrichment of spore-forming Firmicutes at Day 1000 drought: Revisiting Hypothesis 1

Microbes belonging to the phylum Firmicutes can become more abundant during prolonged drought and starvation conditions in soils [26], with supporting evidence from the plant literature [51, 58, 59], due to their metabolic versatility and adaptive strategies (phenotypic heterogeneity/phenotypic plasticity mechanisms) [60]. Firmicutes, particularly those within the classes Bacilli and Clostridia, are known for their ability to form endospores, which allow them to survive in harsh environmental conditions, including drought and nutrient scarcity [61–66]. Endospores enable Firmicutes to survive in a dormant state and become active when conditions improve. These traits make Firmicutes well-suited to thrive in environments with fluctuating nutrient and moisture availability and support the enrichment of classic spore-forming Firmicutes members Bacilli and Clostridia during drought at Day 1000 in our study. Another reason we observe this significant shift in the community towards members of Firmicutes during prolonged drought can be attributed to an enrichment that results from depletion of the viable microbial population, possibly causing the release of microbial biomass and carbon substrates directly benefiting drought-tolerant taxa [17].

The success of endospore-forming Firmicutes in extreme stress environments have been linked to diversified survival strategies and metabolic adaptations with multiple environmental factors favoring their prevalence [67]. The CSR (competitor, stress tolerator, ruderal) framework might apply in this context [68] and explain the dominance of Firmicutes during long-term drought via morphological adaptations (endospores) and stress tolerance features (production of extracellular polymeric substances) to prioritize longevity and efficiency. Eventually, they may outcompete other members of the population while maintaining low levels of activity. Overall, our data support this framework, as we observed a revival of the Firmicutes community when switching from short-term and long-term drought to rewet (copiotrophs/competitors) and an increase in Firmicutes abundance from short-term to prolonged drought (stress tolerators).

Summarizing the above points, we list here three plausible and not mutually exclusive explanations for why we found a higher abundance of certain bacterial taxa at Day 1000 compared to shorter droughts:

1. Stress tolerance: Nutrient depletion/environmental stress allowing certain members to survive via stress tolerance features versus others that die.
2. Habitat differentiation and adaptation: Certain microbes with unique genomic capabilities and functional/morphological adaptations survive as a longer drought period continues; a new environment selecting for species that are adapted to it.
3. A technical artifact: the sequencing depth was not enough to capture certain microbial members in the earlier time points (total mean library size of the samples at Day 30 was 44,130, Day 90 was 32,164, and Day 1000 was 112,719 before rarefaction).

### Differentiating between soil microbial communities from drought and drought-rewet conditions: Revisiting Hypothesis 2

Hypothesis 2 predicted that soils that were dried longer will experience winnowing of the revivable microbial population, and thus the longer-dried soils (especially Day 1000) will show less differentiation in their responses to rewetting. Notably, for the Day 1000 drought samples, the dissimilarity between drought and d+rewet samples was less compared to earlier time points, partially supporting hypothesis 2. However, community size estimates suggest that this reason is likely not due to a winnowed community, since alpha diversity was highest for the Day 1000 drought time point for both the soil depths analyzed (Figure 2B). Instead, the similarity between drought and d+rewet samples appear to be largely driven by increased abundance of Firmicutes at Day 1000 drought and the continued persistence of these taxa and several others post rewetting (Figure 4B).

For the short-term drought treatment, the difference between drought and d+rewet was greater for a few samples, since taxa which were not detected during drought were recruited back to the population upon rewetting, with most of these belonging to the Firmicutes phylum. For some of the other taxa belonging to Proteobacteria, and Verrucomicrobia, there was an opposite response wherein microbes enriched during drought were not persistently detected in the community after rewetting.

However, absolute values of Bray-Curtis distances largely demonstrate an overall dissimilarity in communities between drought and d+rewet for both short-and long-term drought in our study. Interestingly, the relative similarity in microbial community composition observed between long-term drought and subsequent rewet in our study corroborates another study in the literature, though with regards to a short-term drought (28 days) and rewetting for three weeks [69]. This study demonstrated minimal effects on prokaryotic communities, such that only 8% of taxa changed in abundance between drought and d+rewet, with other taxa showing no difference between the treatments [69].

Rewetting soils after a prolonged drought that extends greater than 4 weeks can also cause a ‘lag period’ after which microbial responses show a secondary increase in respiration [70]. This lag period was also reported to increase as drought duration increased and might explain the greater similarity between Day 1000 drought and d+rewet treatments compared to short-term drought. High-intensity perturbations can also abruptly shift the microbial community composition with the persistence of an altered microbiome after soils are rewet to initial moisture status [71]. Finally, the increased abundance of Gram-positive bacteria (Firmicutes and Actinobacteria) during prolonged drought can cause an inherent resistance to drying-rewetting events due to morphological structures such as strong, thick, peptidoglycan cell walls [72].

### Drought duration does not influence the resilience of the community post rewetting: Revisiting Hypothesis 3

A striking observation was that the microbial communities did not recover to pre-drought state post rewetting for both the short-and long-term drought treatments. We expected the communities to recover to pre-drought conditions once samples were rewet following a short-term drought (greater resilience) whereas the expectation for long-term drought was that the communities would be altered significantly to return to a pre-drought condition post rewetting (less resilience) (Figure 6).

There could be several reasons why rewetting may not restore the pre-drought community even during short-term drought. Research has shown that intense “pulse” drought events can significantly reduce microbial community complexity and functionality, with the altered community persisting after soil returns to its previous moisture status [71]. Differential mortality and selection can favor certain stress-tolerant taxa during short-term drought [73] such as Gram-positive Actinobacteria which persist after rewetting. The thick cross-linked peptidoglycan cell walls of Gram-positive bacteria, such as Actinobacteria, act as a physical barrier against osmotic pressure during drying and subsequent rewetting [72, 74].

Resource depletion can also hinder the recovery of a community upon rewetting[75]. While many microbes enter a state of dormancy during dry periods, their resuscitation upon rewetting is not always immediate and may encounter physiological lags that disrupt soil carbon and nitrogen cycling [47, 76]. Finally, as drought alters soil physicochemical properties such as soil moisture, pH, carbon and nitrogen content, a “new” environment may be created, marking a shift in nutrient dynamics that may no longer support the original pre-drought community composition [77].

There are some limitations to our approach and experimental design. First, we chose short-term drought durations for 30 days, and 90 days and long-term drought for 1000 days. Additional time points of sampling incrementing at 6 months would provide a greater temporal resolution of microbial community dynamics throughout the 3-year drought duration. Second, we are unable to provide any insight on the microbial activity or functional dynamics during short and long-term drought, and subsequent reactivation upon rewetting due to lack of metatranscriptomic or metagenomic data. Recent work, however, has shown that mRNA, rRNA, and metabolites can essentially remain frozen during desiccation (dormancy-associated molecular preservation) in certain non-spore forming bacteria, which has challenged RNA based activity assessments in dry environments [78]. However, how these activity patterns pan out during extended drought treatments need further investigation.

Conducting this study in a laboratory setting allowed us to draw new understanding of how soil microorganisms persist in soils, and how they respond to extended drought. We found that microbiome resilience to rewetting after short-term and long-term drought was independent of drought duration, with an altered community (high dissimilarity to pre-drought conditions) after rewetting for all drought durations studied. Surprisingly, the overwhelming presence of certain taxa at Day 1000 drought, which were largely undetected in Day 30, and Day 90 time points, poses new questions about the nature of latent organisms in soils and their role in adaptation and resilience to stress. Future studies aimed at genome-resolved mechanisms of such adaptation will improve our understanding of microbiome resilience to changing weather trajectories and abrupt transitions in soil moisture regime.

## Acknowledgements

We acknowledge Nicholas J. Reichart for conducting the Illumina library prep and sequencing on the MiSeq for a subset of the samples. This research was supported by the U.S. Department of Energy, Office of Science, Biological and Environmental Research as part of the Environmental System Science Program. The Pacific Northwest National Laboratory is operated for DOE by Battelle Memorial Institute under contract DE-AC05-76RL01830. A portion of this research was performed using EMSL, a DOE Office of Science user facility sponsored by the Department of Energy’s Office of Biological and Environmental Research and located at Pacific Northwest National Laboratory.

## Data Availability

Raw DNA sequence data is uploaded to NCBI Sequence Read Archive under project number PRJNA 1450191. Processed data files for analysis are uploaded to ESS-DIVE and can be accessed here https://data.ess-dive.lbl.gov/datasets/doi:10.15485/3376192 [79]. All code used for analysis is uploaded to GitHub (https://github.com/PNNL-TES/TES-drydown.git) and archived at Zenodo (https://doi.org/10.5281/zenodo.20753307) [80].

## Author Contributions

Conceptualization – VLB, SB; Methodology – VLB, SB, KFP, SJF, JZ; Software – SB, KFP, JZ; Validation – SB, KFP; Formal Analysis – SB, KFP; Investigation – SB, KFP, SJF, BBL; Data Curation – SB, KFP, SJF, BBL; Visualization-SB, KFP; Resources – VLB, SJF; Supervision – VLB; Project Administration – VLB, SB; Funding Acquisition – VLB; Writing: original draft – SB; Writing: review, editing – SB, VLB, BBL, KFP, JZ.

