## Supplementary Information for "Drought duration does not impact soil microbiome resilience"

***Corresponding author:**

### Methods

#### 16SrRNA gene amplicon sequencing

PCR reactions were set up as follows to amplify the V4 region of the 16S rRNA gene:

20 uL (for 1X final concentration) of Platinum II 2X Master Mix, 1 uL of forward primer containing index barcode (515f 5'-GTGYCAGCMGCCGCGGTAA-3') (10 uM) for a final concentration of 0.2 uM, 1 uL of reverse primer (806r 5'-GGACTACNVGGGTWTCTAAT-3') (10 uM) for a final concentration of 0.2 uM, 23 uL of nuclease free water, and 5 uL DNA template. PCR program for amplifying target region was set up as follows - 35 cycles of hot start (94 C for 3 minutes), denature (94 C for 45 seconds), anneal (50 C for 60 seconds), extend (72 C for 90 seconds), final extension (72 C for 10 minutes), and hold at 4 C. Picogreen dsDNA quantification assay was used to determine amplicon concentrations. Samples were quantified in triplicate following manufacturer's protocol using 1 uL of DNA template.

*Re-PCR of select samples* Following the Picogreen quantification, several samples were deemed to be below necessary concentration. These samples were amplified again using a lower input of starting material and measured again for amplicon quantification. Unique barcodes were used for the re-amplification of these samples. PCR reactions were set up as follows: 20 uL (for 1X final concentration) of Platinum II 2X Master Mix, 1 uL forward primer containing index barcode (515f 5'-GTGYCAGCMGCCGCGGTAA-3') (10 uM) for a final concentration of 0.2 uM, 1 uL of reverse primer (806r 5'-GGACTACNVGGGTWTCTAAT-3') (10 uM) for a final concentration of 0.2 uM, 26 uL nuclease free water, and 2 uL DNA template. Based on Picogreen quantification results, 200 ng of DNA per sample was pooled. Using the Zymo DNA Clean and Concentrator kit, the pooled sample was cleaned and concentrated into a single tube containing 50 uL of cleaned DNA at a concentration of 5.29 ng/uL. If a sample did not have high enough concentration to pool 200 ng, 40 uL was transferred. The reamplified samples were included in the sample pool instead of the corresponding original PCR product for these samples. Sample was diluted to 2nM and the standard Illumina library prep was followed to load the samples onto the MiSeq for v2 sequencing using a 500-cycle flow cell and a 15 % PhiX spike-in.

### Water extractable organic matter characterization

#### Nuclear magnetic resonance (NMR)

Ten mL of WEOC extract was freeze-dried and then reconstituted with 600 μL of DMSO-$d_6$ into 5 mm NMR tubes and analyzed on a Bruker 9.4 T 400 MHz Avance III NMR spectrometer with a BBFO 5 mm SmartProbe. One-dimensional $^1$H NMR experiments were performed using 1600 scans and a recycle delay of 1 second. Spectral processing included phase and baseline correction, and spectral deconvolution through peak picking by the Global Deconvolution Model in MestreNova v14.1.2-25024 (Mestrelab Research). Peak intensities were normalized to the solvent.

Processing of NMR data was done using the *nmrrr* workflow in R [1]. The spectra were integrated into six regions following [2]: (1) aliphatic polymethylene and methyl groups (0.6–1.3 ppm, referred to as “aliphatic1” here); (2) aliphatic methyl and methylene near O and N (1.3–2.9 ppm, referred to as “aliphatic2” here); (3) O-alkyl, mainly from carbohydrates and lignin (2.9–4.1 ppm); (4) α-proton of peptides (4.1–4.8 ppm); (5) aromatic and phenolic (6.2–7.8 ppm); and (6) amide, from proteins (7.8–8.4 ppm). Integrations in each region were calculated as the sum of all peak areas within the region.

The relative abundance of each region was calculated as the proportion of the signal in each region to the total across all regions, normalized to 100%. As residual water peak (3.33 ppm) occupied most of the O-alkyl region in a non-uniform way across samples, we disregard this region entirely in our analyses. Peaks in the region 2.5-2.75 ppm were also disregarded, to avoid interference of the DMSO-$d_6$ solvent peaks.

**Fourier-transform ion cyclotron resonance mass spectrometry (FT-ICR-MS)**

The WEOC extracts were standardized to 25 ppm C and further purified by solid phase extraction (SPE). Samples were acidified to pH 2 with 85% phosphoric acid and then passed through Bond Elut PPL cartridges (Agilent Technologies) preactivated with $CH_3OH$. The cartridges were washed three times with 10 mM HCl and then dried with $N_2$. The samples were eluted with 1.5 mL $CH_3OH$. The extracts were analyzed at the Environmental Molecular Sciences Laboratory (EMSL), a US-DOE user facility, using a 12T Bruker SolariX FT-ICR-MS spectrometer. 96 individual scans were averaged for each sample and internally calibrated using an organic matter homologous series separated by 14 Da (–CH2 groups). The mass measurement accuracy was less than 1 ppm for singly charged ions across a broad m/z range (m/z 200– 900). Data analysis software (Bruker Daltonik version 4.2) was used to convert raw spectra to a list of m/z values, and absolute intensity threshold set to the default value of 100. Chemical formulae were assigned using the Formularity program, including only peaks with a signal/noise ratio > 7 [3, 4], with the restrictions $C_{1-130}$, $H_{1-200}$, $O_{1-50}$, $N_{0-3}$, $S_{0-3}$, $P_{0-2}$.

Processing of FT-ICR-MS data was done using the *fticrrr* workflow in R [5]. FT-ICR-MS-resolved peaks were analyzed only on a presence/absence basis, due to established issues with intensities and ionization efficiencies in complex matrices using ESI [6]. Only peaks present in 3 or more replicates were considered present within that treatment. The peaks were assigned to one of the following classes following the method of Seidel et al. [7, 8]: (1) polycyclic condensed aromatics ($AI_{mod} > 0.66$); (2) highly aromatic compounds, which include polyphenols and polycyclic aromatic compounds with aliphatic chains ($0.66 > AI_{mod} > 0.50$); (3) highly unsaturated compounds, which include phenols such as soil-derived products of lignin degradation ($AI_{mod}$ > 0.50 and H/C < 1.5); and (4) aliphatic compounds (> H/C ≥ 1.5), including unsaturated aliphatics, N-containing aliphatics, and saturated compounds including fatty and sulfonic acids, and/or carbohydrates. The FT-ICR-MS compound classes are tentative classifications as they are solely based on the indices (O/C and H/C ratios and $AI_{mod}$ values) from the molecular formula, not the molecular structure. Relative abundance values were calculated from count values associated with each observed biomolecule group normalized by the total number of C molecules identified.

### Results

#### Microbial sequence summary

Sequencing was completed as part of a larger dataset containing samples other than those used in the present study. A total of 260 samples were sequenced (253 true samples, 7 negative controls) with 40, 654 features. Out of these, 42 features were identified as contaminants (28 contaminants in WA samples), leaving 40, 612 features for downstream analysis. Data were rarefied at 5000 read depth which removed 43 samples resulting in a total of 210 samples. Missing metadata information for a few samples led to a final sample size of 203 samples. Of these 94 samples were used in the final analysis corresponding to the Washington samples and 27915 taxa in the final Washington count table. Although several samples were removed due to rarefaction, enough replicates were retained for each time point to appropriately conduct statistical analyses.

#### Impact of rarefying reads on indicator taxa pool

To test whether rarefying reads had any contradictory effects on the indicator taxa results by drought duration, we used a differential expression analysis using DeSEq2 without rarefying reads and retaining more samples in our comparisons. This showed that OTUs belonging to Actinobacteriota and Firmicutes were significantly enriched in Day 1000 drought versus Day 30 and Day 90 drought (magnitude of log two-fold change > 2, Figure S3A, B).

**Drought responsive taxa were phylogenetically conserved at higher taxonomic levels**

A majority of the taxa within the top 50 most abundant indicator species pool for drought (Figure 4B) were all within domain Bacteria and represented 11 different phyla (Figure S4A). For the short-term drought, the indicator taxa were mainly within phyla Acidobacteriota, Chloroflexi, Proteobacteria, Verrucomicrobiota, and Methylomirabilota, whereas, for the long-term (Day 1000) drought indicator taxa, responses were conserved within three core phyla: Myxococcota, Firmicutes and Actinobacteriota (Figure S4B).

### Supplementary Tables

**Table S1.** Soil characteristics, as reported in [9].

| | |
|---|---|
| Total C (%) | 10.32 ± 0.45 |
| Total N (%) | 0.59 ± 0.01 |
| Total organic C (%) | 9.94 ± 0.47 |
| pH | 5.82 ± 0.05 |
| Sand (%) | 14 ± 2.12 |
| Silt (%) | 58.5 ± 1.5 |
| Clay (%) | 27.75 ± 1.65 |
| Texture | silty clay loam |

**Table S2:** PERMANOVA statistics for microbial communities as they differ by saturation, time, soil depth, and drying method. All statistics are based on Bray-Curtis distances between samples.

| Factor | Degrees of Freedom | Sum of Squares | $R^2$ | F value | P value |
|---|---|---|---|---|---|
| **saturation** | 2 | 6.609 | 0.19301 | 13.9831 | 0.001 |
| **time** | 2 | 3.595 | 0.10498 | 7.6058 | 0.001 |
| **depth** | 1 | 0.8 | 0.02336 | 3.3849 | 0.001 |
| **drying** | 1 | 0.276 | 0.00807 | 1.1693 | 0.26 |
| **saturation:time** | 2 | 2.584 | 0.07546 | 5.4667 | 0.001 |
| **saturation:depth** | 2 | 0.716 | 0.02091 | 1.5147 | 0.045 |
| **time:depth** | 2 | 0.726 | 0.02119 | 1.5351 | 0.034 |
| **saturation:drying** | 1 | 0.304 | 0.00889 | 1.2884 | 0.159 |
| **time:drying** | 2 | 0.402 | 0.01175 | 0.8513 | 0.714 |
| **depth:drying** | 1 | 0.164 | 0.00478 | 0.6929 | 0.812 |
| **saturation:time:depth** | 2 | 0.588 | 0.01718 | 1.2446 | 0.178 |
| **saturation:time:drying** | 2 | 0.464 | 0.01356 | 0.9825 | 0.456 |
| **saturation:depth:drying** | 1 | 0.191 | 0.00558 | 0.8088 | 0.68 |
| **time:depth:drying** | 2 | 0.359 | 0.01047 | 0.7588 | 0.863 |
| **saturation:time:depth:drying** | 2 | 0.393 | 0.01149 | 0.8323 | 0.705 |
| **Residual** | 68 | 16.07 | 0.46931 | | |
| **Total** | 93 | 34.242 | 1 | | |

**Table S3:** Indicator taxa by drought duration.

| OTU | Day 1000 | Day 30 | Day 90 | Day 0 | index | stat | p.value | Phylum | Class | Order | Family | Genus |
|---|---|---|---|---|---|---|---|---|---|---|---|---|
| 180a8b3ab79f7c154 15c90150a19883d | 1 | 0 | 0 | 0 | 1 | 0.9289697 | 0.01504651 | p__Firmicutes | c__Bacilli | o__Paenibacillales | f__Paenibacillaceae | g__Paenibacillus |
| 26005dfb7ad2e8cdb b139846af329663 | 1 | 0 | 0 | 0 | 1 | 0.92494985 | 0.01504651 | p__Firmicutes | c__Bacilli | o__Bacillales | f__Bacillaceae | g__Bacillus |
| 88a607b6dfeddc8fb5 b89bc2ce51623a | 0 | 0 | 0 | 1 | 4 | 0.90287648 | 0.01504651 | p__Bacteroidota | c__Bacteroidia | o__Flavobacteriales | f__Flavobacteriaceae | g__Flavobacterium |
| 9847d2b1b764509d8 fb3ec9fa2726d6c | 0 | 0 | 0 | 1 | 4 | 0.86785653 | 0.01504651 | p__Proteobacteria | c__Gammaproteob acteria | o__Burkholderiales | f__SC-I-84 | g__SC-I-84 |
| 33d917f857a921756 e518b891371da1e | 1 | 0 | 0 | 0 | 1 | 0.85723777 | 0.01504651 | p__Myxococcota | c__Polyangia | o__Haliangiales | f__Haliangiaceae | g__Haliangium |
| d36957a0d3172d2e5 612266f417bbed6 | 0 | 0 | 0 | 1 | 4 | 0.84926794 | 0.01504651 | p__Proteobacteria | c__Gammaproteob acteria | o__Burkholderiales | f__Nitrosomonadace ae | g__MND1 |
| 88fb15f05df0863489 63d416fa3f3077 | 0 | 0 | 0 | 1 | 4 | 0.84756099 | 0.01504651 | p__Acidobacteriota | c__Blastocatellia | o__Blastocatellales | f__Blastocatellaceae | g__JGI_0001001-H03 |
| e4ee05bbf4ea42bb7 404e560a5998628 | 0 | 0 | 0 | 1 | 4 | 0.84689862 | 0.01504651 | p__Bacteroidota | c__Bacteroidia | o__Chitinophagales | f__Chitinophagaceae | g__Puia |
| 5958fd2b7ec972907 e061e8f4d0c49b8 | 0 | 0 | 0 | 1 | 4 | 0.84673345 | 0.01504651 | p__Acidobacteriota | c__Subgroup_22 | o__Subgroup_22 | f__Subgroup_22 | g__Subgroup_22 |
| 762facf339b485a7d8 2a2f3fdef5d772 | 0 | 0 | 0 | 1 | 4 | 0.84642433 | 0.01504651 | p__Nitrospirota | c__Nitrospiria | o__Nitrospirales | f__Nitrospiraceae | g__Nitrospira |
| 8cff7bc3fb5f7586f2 6e4837583cb108 | 0 | 0 | 0 | 1 | 4 | 0.8446811 | 0.01504651 | p__Proteobacteria | c__Gammaproteob acteria | o__Cellvibrionales | f__Halieaceae | g__OM60(NOR5)_cl ade |
| 25e9838ddd33c0994 e7d6f6992e39de7 | 1 | 0 | 0 | 0 | 1 | 0.84058112 | 0.01504651 | p__Firmicutes | c__Bacilli | o__Paenibacillales | f__Paenibacillaceae | g__Cohnella |
| ed4eb0c93690ed93b 1306bbc507ef2ba | 0 | 0 | 0 | 1 | 4 | 0.83149384 | 0.01504651 | p__Proteobacteria | c__Gammaproteob acteria | o__Burkholderiales | f__Nitrosomonadace ae | g__mle1-7 |

| | | | | | | | | | | | | |
|---|---|---|---|---|---|---|---|---|---|---|---|---|
| a1b2e6767a5ecfa91b0843b785a3e44c | 0 | 0 | 0 | 1 | 4 | 0.82967582 | 0.01504651 | p__Bacteroidota | c__Bacteroidia | o__Chitinophagales | f__Chitinophagaceae | g__Terrimonas |
| e4d87db692a9367083ee3831320420a2 | 1 | 0 | 0 | 0 | 1 | 0.82885948 | 0.01504651 | p__Bdellovibrionota | c__Oligoflexia | o__0319-6G20 | f__0319-6G20 | g__0319-6G20 |
| a8835f7d0e717b389f66bf0ea1cf4718 | 0 | 0 | 0 | 1 | 4 | 0.82556388 | 0.01504651 | p__Latescibacterota | c__Latescibacterota | o__Latescibacterota | f__Latescibacterota | g__Latescibacterota |
| 5b2529faa44a3679b887ecd91b6b4149 | 0 | 1 | 1 | 0 | 8 | 0.82053583 | 0.01504651 | p__Chloroflexi | c__KD4-96 | o__KD4-96 | f__KD4-96 | g__KD4-96 |
| 4804669ff00c7ec682696fdd03d60923 | 0 | 0 | 0 | 1 | 4 | 0.80370004 | 0.01504651 | p__Proteobacteria | c__Gammaproteobacteria | o__Xanthomonadales | f__Xanthomonadaceae | g__Arenimonas |
| e3b25be4febe41497c6bd9a899eb32aa | 1 | 0 | 0 | 0 | 1 | 0.80128177 | 0.01504651 | p__Myxococcota | c__Myxococcia | o__Myxococcales | f__Myxococcaceae | g__P3OB-42 |
| 120eba657e42a11a5c29f97b90f02035 | 1 | 0 | 0 | 0 | 1 | 0.79852482 | 0.01504651 | p__Actinobacteriota | c__Actinobacteria | o__Streptomycetales | f__Streptomycetaceae | g__Streptomyces |
| f073df90e41bdd0c9674d8de667f5f32 | 0 | 0 | 0 | 1 | 4 | 0.79443368 | 0.01504651 | p__Proteobacteria | c__Alphaproteobacteria | o__Caulobacterales | f__Hyphomonadaceae | g__SWB02 |
| 8206db4b8b8bbd572d04ab96cb720f52 | 1 | 0 | 0 | 0 | 1 | 0.79332689 | 0.01504651 | p__Actinobacteriota | c__Thermoleophilia | o__Solirubrobacterales | f__67-14 | g__67-14 |
| 6be01b304ffd87213eb490e18acdcec4 | 1 | 0 | 0 | 0 | 1 | 0.7910387 | 0.01504651 | p__Actinobacteriota | c__Thermoleophilia | o__Solirubrobacterales | f__Solirubrobacteraceae | g__Solirubrobacter |
| d20b46e3c9d79a8e49a48f112fc03d4f | 1 | 0 | 0 | 0 | 1 | 0.77894184 | 0.01504651 | p__Firmicutes | c__Clostridia | o__Peptostreptococcales-Tissierellales | f__Peptostreptococcaceae | g__Romboutsia |
| a2720964e08520e61a2ae03ffb04f083 | 0 | 0 | 0 | 1 | 4 | 0.7765575 | 0.01504651 | p__Latescibacterota | c__Latescibacteria | o__Latescibacterales | f__Latescibacteraceae | g__Latescibacteraceae |
| 40fa96a2f7fc7b939f5eb0d52d7e5b3c | 0 | 0 | 0 | 1 | 4 | 0.76829119 | 0.01504651 | p__Proteobacteria | c__Gammaproteobacteria | o__Burkholderiales | f__Nitrosomonadaceae | g__IS-44 |
| d7ce932b3b7fd20cc7a13918ee716f0a | 1 | 0 | 0 | 0 | 1 | 0.76309283 | 0.01504651 | p__Firmicutes | c__Desulfitobacteriia | o__Desulfitobacteriales | f__Desulfitobacteriaceae | g__Desulfosporosinus |
| 18c95fb5b9dbfd87d20f1dcdc8488cba | 1 | 0 | 0 | 0 | 1 | 0.75951808 | 0.02121311 | p__Firmicutes | c__Clostridia | o__Clostridiales | f__Caloramatoraceae | g__Fonticella |

| | | | | | | | | | | | | |
|---|---|---|---|---|---|---|---|---|---|---|---|---|
| 23bc097ab0bec1cc29d67480c567fb6b | 0 | 0 | 0 | 1 | 4 | 0.75309706 | 0.02121311 | p__Proteobacteria | c__Alphaproteobacteria | o__Rhizobiales | f__A0839 | g__A0839 |
| aff33135a79f13e6c4ef2dd338bd14c5 | 0 | 0 | 0 | 1 | 4 | 0.75180507 | 0.01504651 | p__Planctomycetota | c__OM190 | o__OM190 | f__OM190 | g__OM190 |
| f4ff7f76850896c5c1bcd6931261882c | 0 | 1 | 1 | 0 | 8 | 0.74961441 | 0.01504651 | p__Verrucomicrobiota | c__Verrucomicrobiae | o__S-BQ2-57_soil_group | f__S-BQ2-57_soil_group | g__S-BQ2-57_soil_group |
| d3b52387360ceed8f554547f02b16f85 | 1 | 0 | 0 | 0 | 1 | 0.74815907 | 0.02695833 | p__Actinobacteriota | c__Actinobacteria | o__Micromonosporales | f__Micromonosporaceae | g__Micromonospora |
| 3807a2439c4c8b2085b7dd9c9e995428 | 0 | 0 | 0 | 1 | 4 | 0.74545343 | 0.02121311 | p__Proteobacteria | c__Gammaproteobacteria | o__Salinisphaerales | f__Solimonadaceae | g__Polycyclovorans |
| f6da163672ad774ddc3ca4bc4dc96245 | 0 | 0 | 0 | 1 | 4 | 0.74464745 | 0.01504651 | p__Proteobacteria | c__Gammaproteobacteria | o__KI89A_clade | f__KI89A_clade | g__KI89A_clade |
| c27eddb9cd73ca71dcddf8a8d16bcb2b | 1 | 0 | 0 | 0 | 1 | 0.73100793 | 0.02121311 | p__Firmicutes | c__Clostridia | o__Clostridia | f__Hungateiclostridiaceae | g__Ruminiclostridium |
| 4fcfe434053e932f99d83b2c48bea73c | 0 | 0 | 0 | 1 | 4 | 0.73010054 | 0.01504651 | p__Bacteroidota | c__Bacteroidia | o__Sphingobacteriales | f__AKYH767 | g__AKYH767 |
| 811ef0f05f1341efae2dabdf27a7b79a | 0 | 1 | 1 | 1 | 14 | 0.72993664 | 0.01504651 | p__Acidobacteriota | c__Vicinamibacteria | o__Subgroup_17 | f__Subgroup_17 | g__Subgroup_17 |
| ffa811c78b86c1935fb127fa54f4b756 | 1 | 0 | 0 | 0 | 1 | 0.72815657 | 0.02121311 | p__Myxococcota | c__Polyangia | o__Polyangiales | f__Polyangiaceae | g__Sorangium |
| 0af3a1da0bc47c96a1700ebd61f39269 | 1 | 0 | 0 | 0 | 1 | 0.72683806 | 0.02695833 | p__Firmicutes | c__Bacilli | o__Alicyclobacillales | f__Alicyclobacillaceae | g__Tumebacillus |
| 594fb62b72cd8f8ebab8e7173fb7340a | 1 | 0 | 0 | 0 | 1 | 0.72276418 | 0.02121311 | p__Actinobacteriota | c__Actinobacteria | o__Frankiales | f__Frankiaceae | g__Frankia |
| 2969ec24415ae1ed31f6260afa476743 | 1 | 0 | 0 | 0 | 1 | 0.72176538 | 0.01504651 | p__Actinobacteriota | c__Thermoleophilia | o__Solirubrobacterales | f__Solirubrobacteraceae | g__Conexibacter |
| 41c0870e909a9031905df5780affefef | 0 | 0 | 0 | 1 | 4 | 0.71989105 | 0.01504651 | p__Proteobacteria | c__Alphaproteobacteria | o__Paracaedibacterales | f__Paracaedibacteraceae | g__Candidatus_Paracaedibacter |
| d1040bdca6a891ebd5fc8dc0e8a9b767 | 1 | 0 | 0 | 0 | 1 | 0.71811104 | 0.01504651 | p__Firmicutes | c__Clostridia | o__Clostridia_vadinBB60_group | f__Clostridia_vadinBB60_group | g__Clostridia_vadinBB60_group |

| | | | | | | | | | | | | |
|---|---|---|---|---|---|---|---|---|---|---|---|---|
| bde251162ad0cea92c6d0dfacda4ac10 | 0 | 0 | 0 | 1 | 4 | 0.70866466 | 0.01504651 | p__Proteobacteria | c__Gammaproteobacteria | o__KF-JG30-C25 | f__KF-JG30-C25 | g__KF-JG30-C25 |
| 2573d88484d9b2979456e4a5accbabc6 | 0 | 1 | 1 | 1 | 14 | 0.70662432 | 0.02695833 | p__Acidobacteriota | c__Acidobacteriae | o__Bryobacterales | f__Bryobacteraceae | g__Bryobacter |
| 0952299d87618319b70016986c74ab69 | 1 | 0 | 0 | 0 | 1 | 0.70424093 | 0.02121311 | p__Myxococcota | c__Polyangia | o__Polyangiales | f__Polyangiaceae | g__Pajaroellobacter |
| ade04fd51fe2a7ea6a5de7be6c690573 | 0 | 0 | 0 | 1 | 4 | 0.70053421 | 0.02121311 | p__Proteobacteria | c__Gammaproteobacteria | o__JG36-TzT-191 | f__JG36-TzT-191 | g__JG36-TzT-191 |
| 2e4a2e5ad75f2d321aadefddacb2fca7 | 0 | 0 | 0 | 1 | 4 | 0.69912117 | 0.01504651 | p__Proteobacteria | c__Gammaproteobacteria | o__Gammaproteobacteria_Incertae_Sedis | f__Unknown_Family | g__Acidibacter |
| 1057558de242812fdb138341e65f73d0 | 0 | 0 | 0 | 1 | 4 | 0.69504472 | 0.01504651 | p__NB1-j | c__NB1-j | o__NB1-j | f__NB1-j | g__NB1-j |
| 16ab9298f3d7f93f8ac8598c124cbd18 | 0 | 1 | 1 | 1 | 14 | 0.6944908 | 0.02121311 | p__Acidobacteriota | c__Blastocatellia | o__Pyrinomonadales | f__Pyrinomonadaceae | g__RB41 |
| ee47186ecd61e60519bbc886513b03de | 1 | 0 | 0 | 0 | 1 | 0.69430987 | 0.02695833 | p__Firmicutes | c__Clostridia | o__Lachnospirales | f__Lachnospiraceae | g__Anaerocolumna |
| 20ff995601305d0e5b1a189493d2f863 | 1 | 0 | 0 | 0 | 1 | 0.69385327 | 0.01504651 | p__Firmicutes | c__Clostridia | o__Thermincolales | f__Thermincolaceae | g__Thermincola |
| 3d8fb6f4ed7962d88c2f3ae9411d13de | 1 | 0 | 0 | 0 | 1 | 0.69378191 | 0.02121311 | p__Firmicutes | c__Bacilli | o__Aneurinibacillales | f__Aneurinibacillaceae | g__Aneurinibacillus |
| c01565ffe4b71405677739dcbff6041e | 0 | 0 | 0 | 1 | 4 | 0.69346431 | 0.02695833 | p__Bacteroidota | c__Bacteroidia | o__Chitinophagales | f__Chitinophagaceae | g__Ferruginibacter |
| ed162f239f7757cb436d4527f9b3b892 | 0 | 1 | 1 | 1 | 14 | 0.69239742 | 0.02121311 | p__Acidobacteriota | c__Blastocatellia | o__11-24 | f__11-24 | g__11-24 |
| cf20d90736ccd5364db38c3868a48c44 | 0 | 0 | 0 | 1 | 4 | 0.69138556 | 0.03478495 | p__Proteobacteria | c__Gammaproteobacteria | o__WD260 | f__WD260 | g__WD260 |
| 24d50d3c31d64462240e330c879c41e7 | 1 | 0 | 0 | 0 | 1 | 0.68489623 | 0.02121311 | p__Firmicutes | c__Clostridia | o__Clostridiales | f__Clostridiaceae | g__Clostridium_sensu_stricto_12 |
| 858603b5b6c7332924623a52faa10212 | 1 | 0 | 0 | 0 | 1 | 0.68266766 | 0.02695833 | p__Firmicutes | c__Clostridia | o__Clostridiales | f__Clostridiaceae | g__Clostridium_sensu_stricto_1 |

| | | | | | | | | | | | | |
|---|---|---|---|---|---|---|---|---|---|---|---|---|
| 2f031a61a320cf2abf5952f3deedcb36 | 0 | 1 | 1 | 0 | 8 | 0.68265047 | 0.02695833 | p__Chloroflexi | c__Dehalococcoidia | o__S085 | f__S085 | g__S085 |
| 571da70e16365f2cc78b95f01af868be | 0 | 1 | 1 | 1 | 14 | 0.68036101 | 0.01504651 | p__Verrucomicrobiota | c__Verrucomicrobiae | o__Pedosphaerales | f__Pedosphaeraceae | g__Pedosphaeraceae |
| ede15fb563222a59b54464ff3857badb | 0 | 0 | 0 | 1 | 4 | 0.67814418 | 0.02121311 | p__Proteobacteria | c__Gammaproteobacteria | o__PLTA13 | f__PLTA13 | g__PLTA13 |
| a8dc5aeb11a8e3f8ac305f73ed6c240e | 1 | 0 | 0 | 0 | 1 | 0.67582564 | 0.02695833 | p__Firmicutes | c__Bacilli | o__Bacillales | f__Planococcaceae | g__Lysinibacillus |
| 7c6f15b7e3c29be06d538c984f1a5f9e | 0 | 1 | 1 | 0 | 8 | 0.67506898 | 0.02695833 | p__Chloroflexi | c__Anaerolineae | o__SBR1031 | f__SBR1031 | g__SBR1031 |
| 45351e42da480f7340ccd0703729b83c | 0 | 0 | 0 | 1 | 4 | 0.67350609 | 0.03478495 | p__Bacteroidota | c__Bacteroidia | o__Cytophagales | f__Hymenobacteraceae | g__Adhaeribacter |
| 3a66a38841ba6877568198b9dac3a1dc | 0 | 0 | 0 | 1 | 4 | 0.67290736 | 0.01504651 | p__Proteobacteria | c__Gammaproteobacteria | o__Xanthomonadales | f__Xanthomonadaceae | g__Lysobacter |
| 7c967aff27178682f0d793ac890c2420 | 0 | 1 | 1 | 0 | 8 | 0.6632582 | 0.02121311 | p__Actinobacteriota | c__Acidimicrobiia | o__IMCC26256 | f__IMCC26256 | g__IMCC26256 |
| 0cc1019e6265a3f911ad824615461786 | 0 | 0 | 0 | 1 | 4 | 0.66132494 | 0.02695833 | p__Proteobacteria | c__Alphaproteobacteria | o__Sphingomonadales | f__Sphingomonadaceae | g__Sphingomonas |
| fbfbda0e90e9252d0f7469dc34ce18a0 | 1 | 0 | 0 | 0 | 1 | 0.65954295 | 0.01504651 | p__Firmicutes | c__Limnochordia | o__Limnochordia | f__Limnochordia | g__Hydrogenispora |
| 35d20da789dde8d76e4389fd6d30cc9d | 1 | 0 | 0 | 0 | 1 | 0.65414913 | 0.02121311 | p__Actinobacteriota | c__Actinobacteria | o__Micromonosporales | f__Micromonosporaceae | g__Dactylosporangium |
| 313a85f98077c1aa5b5ab76f7fb4b64a | 0 | 0 | 0 | 1 | 4 | 0.6539046 | 0.03478495 | p__Proteobacteria | c__Gammaproteobacteria | o__Burkholderiales | f__Nitrosomonadaceae | g__Ellin6067 |
| 29f5776f742a007c913db3bececbba42 | 0 | 1 | 1 | 1 | 14 | 0.65348399 | 0.03044706 | p__Acidobacteriota | c__Vicinamibacteria | o__Vicinamibacterales | f__Vicinamibacteraceae | g__Vicinamibacteraceae |
| 486e8a1c521677e7cc6941dee898e39b | 1 | 0 | 0 | 0 | 1 | 0.64908563 | 0.03044706 | p__Firmicutes | c__Bacilli | o__Bacillales | f__Bacillaceae | g__Fictibacillus |
| 191fb83796ae38927791b7d1e1f55060 | 1 | 0 | 0 | 0 | 1 | 0.64869407 | 0.04129787 | p__Firmicutes | c__Bacilli | o__Alicyclobacillales | f__Alicyclobacillaceae | g__Alicyclobacillus |

| | | | | | | | | | | | | |
|---|---|---|---|---|---|---|---|---|---|---|---|---|
| c9f7e7b1bbc0cd09a24c221820329012 | 1 | 0 | 0 | 0 | 1 | 0.64833287 | 0.03478495 | p__Myxococcota | c__Polyangia | o__mle1-27 | f__mle1-27 | g__mle1-27 |
| 6626f9c2e3859382d416f81cb71595bb | 1 | 0 | 0 | 0 | 1 | 0.6464548 | 0.03044706 | p__Firmicutes | c__Negativicutes | o__Veillonellales-Selenomonadales | f__Sporomusaceae | g__Pelosinus |
| b689455693f0c3901ebcbe9ada2a745f | 0 | 1 | 1 | 1 | 14 | 0.64419615 | 0.03044706 | p__Planctomycetota | c__Pla4_lineage | o__Pla4_lineage | f__Pla4_lineage | g__Pla4_lineage |
| e7e704c8cee3aae27ceccf46fd75d332 | 1 | 0 | 0 | 0 | 1 | 0.64176341 | 0.03044706 | p__Firmicutes | c__Desulfitobacteriia | o__Desulfitobacteriales | f__Desulfitobacteriaceae | g__Desulfitobacterium |
| bf0515a90c711833f7adddd9c8177cd | 0 | 1 | 1 | 1 | 14 | 0.63725094 | 0.04484158 | p__Acidobacteriota | c__Subgroup_25 | o__Subgroup_25 | f__Subgroup_25 | g__Subgroup_25 |
| bc7a67999cc192f4c4a021880636ef97 | 1 | 0 | 0 | 0 | 1 | 0.63571427 | 0.02121311 | p__Firmicutes | c__Clostridia | o__Oscillospirales | f__Ruminococcaceae | g__Candidatus_Soleaferrea |
| 5b2fed21a9ae1266d7630fbd98639f48 | 0 | 1 | 1 | 1 | 14 | 0.63493071 | 0.04484158 | p__RCP2-54 | c__RCP2-54 | o__RCP2-54 | f__RCP2-54 | g__RCP2-54 |
| 82a260faafef55efd1b8176ecb781ecf | 1 | 0 | 0 | 0 | 1 | 0.63125721 | 0.04837383 | p__Firmicutes | c__Clostridia | o__Clostridiales | f__Clostridiaceae | g__Clostridium_sensu_stricto_9 |
| 3d7ea4dffab1130cdc0e3659dae0c1ef | 0 | 1 | 1 | 1 | 14 | 0.63065064 | 0.01504651 | p__Methylomirabilota | c__Methylomirabilia | o__Rokubacteriales | f__WX65 | g__WX65 |
| 0f700b983207ec336ed4b4679028b514 | 0 | 1 | 1 | 1 | 14 | 0.62853225 | 0.04484158 | p__Methylomirabilota | c__Methylomirabilia | o__Rokubacteriales | f__Rokubacteriales | g__Rokubacteriales |
| e5e1c77c34ad7f331634811f5de60b32 | 0 | 1 | 1 | 1 | 14 | 0.62846134 | 0.02695833 | p__Proteobacteria | c__Alphaproteobacteria | o__Rhizobiales | f__KF-JG30-B3 | g__KF-JG30-B3 |
| 5c6a9e78d6c0cddb7bcc40b510d9e383 | 1 | 0 | 0 | 0 | 1 | 0.6238686 | 0.03044706 | p__Myxococcota | c__Polyangia | o__Polyangiales | f__Polyangiaceae | g__Minicystis |
| 4cc25ba756f687276764f08c1ebdceb1 | 0 | 0 | 0 | 1 | 4 | 0.62253875 | 0.03044706 | p__Planctomycetota | c__Phycisphaerae | o__Tepidisphaerales | f__WD2101_soil_group | g__WD2101_soil_group |
| f8b8173289056f3442317b4669562a60 | 0 | 1 | 1 | 1 | 14 | 0.61791274 | 0.04484158 | p__Proteobacteria | c__Gammaproteobacteria | o__Burkholderiales | f__TRA3-20 | g__TRA3-20 |
| a3b6cbaee0bed371cc173fede7afb44b | 1 | 1 | 1 | 0 | 11 | 0.61753046 | 0.02121311 | p__Actinobacteriota | c__MB-A2-108 | o__MB-A2-108 | f__MB-A2-108 | g__MB-A2-108 |

| | | | | | | | | | | | | |
|---|---|---|---|---|---|---|---|---|---|---|---|---|
| b48a188a24ed126894db7d73bd6617cc | 1 | 1 | 1 | 0 | 11 | 0.61748573 | 0.03478495 | p__Actinobacteriota | c__Thermoleophilia | o__Gaiellales | f__Gaiellaceae | g__Gaiella |
| a2fdd8fd2907924c8bb04ebda7ac9cc7 | 1 | 0 | 0 | 0 | 1 | 0.61497903 | 0.03044706 | p__Firmicutes | c__Clostridia | o__Oscillospirales | f__Oscillospiraceae | g__Sporobacter |
| 0f4827ad5efbc2cb4863346c87936238 | 0 | 0 | 1 | 1 | 10 | 0.61440174 | 0.03044706 | p__Planctomycetota | c__Planctomycetes | o__Pirellulales | f__Pirellulaceae | g__Pirellula |
| 009a9155135dd33d7cd0abb7258261e2 | 0 | 0 | 0 | 1 | 4 | 0.60951004 | 0.03044706 | p__Bacteroidota | c__Bacteroidia | o__Chitinophagales | f__Chitinophagaceae | g__Dinghuibacter |
| 432b01066dfe0725b4830272bff0caff | 1 | 0 | 0 | 0 | 1 | 0.60663774 | 0.04484158 | p__Firmicutes | c__Bacilli | o__Thermoactinomycetales | f__Thermoactinomycetaceae | g__Thermoactinomyces |
| 25da1a2223a07f6386538f347b59b16b | 1 | 0 | 0 | 0 | 1 | 0.60301401 | 0.04484158 | p__Bacteroidota | c__Bacteroidia | o__Cytophagales | f__Cytophagaceae | g__Sporocytophaga |
| 41b5dae7377b7098ee5e3c8ed7dc60f9 | 1 | 0 | 0 | 0 | 1 | 0.60182838 | 0.03044706 | p__Firmicutes | c__Bacilli | o__Thermoactinomycetales | f__Thermoactinomycetaceae | g__Lihuaxuella |
| 13801e7bc71636162780bd7b6d06bf65 | 1 | 0 | 0 | 0 | 1 | 0.59783252 | 0.03044706 | p__Myxococcota | c__Polyangia | o__Polyangiales | f__Phaselicystidaceae | g__Phaselicystis |
| a09c0c4f5bfda19fb24ab7decd839d32 | 1 | 0 | 0 | 0 | 1 | 0.59608301 | 0.04837383 | p__Firmicutes | c__Bacilli | o__Thermoactinomycetales | f__Thermoactinomycetaceae | g__Thermoactinomycetaceae |
| de0954e4f3fc0f3a550fe89a56fd4560 | 1 | 0 | 0 | 0 | 1 | 0.59589438 | 0.03478495 | p__Firmicutes | c__Clostridia | o__Oscillospirales | f__Ethanoligenenaceae | g__Incertae_Sedis |
| 6e4b35aa6722d7614f574250954cfb8a | 1 | 0 | 0 | 0 | 1 | 0.59509988 | 0.03478495 | p__Firmicutes | c__Bacilli | o__Thermoactinomycetales | f__Thermoactinomycetaceae | g__Geothermomicrobium |
| 35f23ca925e88b47f5d3ec9f1c689432 | 1 | 0 | 0 | 0 | 1 | 0.58749666 | 0.03044706 | p__Actinobacteriota | c__Acidimicrobiia | o__Microtrichales | f__Ilumatobacteraceae | g__Ilumatobacter |
| c63a29416a7eabd7ff67a49489118771 | 0 | 1 | 1 | 0 | 8 | 0.58514455 | 0.04837383 | p__Proteobacteria | c__Alphaproteobacteria | o__Rhizobiales | f__Xanthobacteraceae | g__Pseudolabrys |
| 4a7dd9b8e75b4b720d8aba0c7d1ba082 | 1 | 0 | 0 | 0 | 1 | 0.57615214 | 0.04837383 | p__Verrucomicrobiota | c__Chlamydiae | o__Chlamydiales | f__Parachlamydiaceae | g__Candidatus_Protochlamydia |
| c62774eb8230ddbdc937c240c25753e5 | 1 | 0 | 0 | 0 | 1 | 0.56839856 | 0.04837383 | p__Firmicutes | c__Clostridia | o__Oscillospirales | f__Ruminococcaceae | g__Ruminococcus |

| | | | | | | | | | | | | |
|---|---|---|---|---|---|---|---|---|---|---|---|---|
| 7e3162f0b727bb7d322b493f5dda9a8a | 0 | 1 | 1 | 0 | 8 | 0.56602005 | 0.04484158 | p__Acidobacteriota | c__Thermoanaerobaculia | o__Thermoanaerobaculales | f__Thermoanaerobaculaceae | g__Subgroup_10 |
| f004908a60fef5ccc0caa58fd4cbbca7 | 1 | 0 | 0 | 0 | 1 | 0.55854322 | 0.04837383 | p__Firmicutes | c__Clostridia | o__Lachnospirales | f__Lachnospiraceae | g__Tyzzerella |
| 69f502de8061943a80a1e0233fcea621 | 0 | 0 | 0 | 1 | 4 | 0.54977054 | 0.03478495 | p__Proteobacteria | c__Gammaproteobacteria | o__Steroidobacterales | f__Woeseiaceae | g__JTB255_marine_benthic_group |
| 547232d63dba7161add4d20bed279403 | 0 | 0 | 0 | 1 | 4 | 0.53398872 | 0.02121311 | p__Verrucomicrobiota | c__Chlamydiae | o__Chlamydiales | f__cvE6 | g__cvE6 |

**Table S4:** PERMANOVA statistics for microbial communities as they differ by saturation levels, separated by drought duration and soil depth A. Without upland samples, B. With upland samples

**A.**

| **Treatment** | **Factor (levels)** | **$R^2$** | **F value** | **P value** |
|---|---|---|---|---|
| 30d (0-5 cm) | Saturation (drought v. d+rewet) | 0.43 | 7.46 | 0.002 |
| 90d (0-5 cm) | | 0.43 | 3.76 | 0.04 |
| **1000d (0-5 cm)** | | **0.47** | **8.02** | **<0.001** |
| 30d (5cm-end) | | 0.25 | 3.04 | 0.002 |
| 90d (5cm-end) | | 0.50 | 5.98 | 0.02 |
| **1000d (5cm-end)** | | **0.55** | **10.83** | **<0.001** |

**B.**

| **Treatment** | **Factor (levels)** | **$R^2$** | **F value** | **P value** |
|---|---|---|---|---|
| 30d (0-5 cm) | Saturation (drought v. d+rewet) | 0.25 | 5.1 | <0.001 |
| 90d (0-5 cm) | | 0.34 | 4.7 | 0.005 |
| **1000d (0-5 cm)** | | **0.41** | **9.54** | **<0.001** |
| 30d (5cm-end) | | 0.24 | 4.5 | <0.001 |
| 90d (5cm-end) | | 0.34 | 6.24 | 0.005 |

| | | | | |
|---|---|---|---|---|
| **1000d (5cm-end)** | | **0.47** | **12.51** | **<0.001** |

**Table S5:** PERMANOVA statistics for FTICR A. for both 0-5 cm depth and 5cm-end depths B. including top depth only

**A.**

| Factor | Degrees of Freedom | Sum of Squares | $R^2$ | F value | P value |
|---|---|---|---|---|---|
| depth | 1 | 0.0506674 | 0.1149601 | 88.057141 | 0.001 |
| time | 2 | 0.2056311 | 0.4665603 | 178.687892 | 0.001 |
| saturation | 1 | 0.1540093 | 0.3494347 | 267.659890 | 0.001 |
| drying | 1 | 0.0018533 | 0.0042049 | 3.220881 | 0.067 |
| Residual | 89 | 0.0512099 | 0.1161911 | NA | NA |
| Total | 94 | 0.4407384 | 1.0000000 | NA | NA |

**B.**

| Factor | Degrees of Freedom | Sum of Squares | $R^2$ | F value | P value |
|---|---|---|---|---|---|
| time | 2 | 0.1219450 | 0.7254832 | 189.9259569 | 0.001 |
| saturation | 1 | 0.0420732 | 0.2503049 | 131.0558189 | 0.001 |
| drying | 1 | 0.0000653 | 0.0003882 | 0.2032546 | 0.661 |
| Residual | 43 | 0.0138044 | 0.0821262 | NA | NA |
| Total | 47 | 0.1680880 | 1.0000000 | NA | NA |

**Table S6:** PERMANOVA statistics for NMR.

| term | Degrees of Freedom | Sum of Squares | $R^2$ | F value | P value |
|---|---|---|---|---|---|
| time | 2 | 0.7142123 | 0.1163433 | 3.7384819 | 0.009 |
| saturation | 1 | 2.4083480 | 0.3923136 | 25.2125733 | 0.001 |
| drying | 1 | -0.0286895 | -0.0046734 | -0.3003454 | 1.000 |
| Residual | 31 | 2.9611729 | 0.4823673 | NA | NA |
| Total | 36 | 6.1388341 | 1.0000000 | NA | NA |

### Supplementary Figures

**A.**

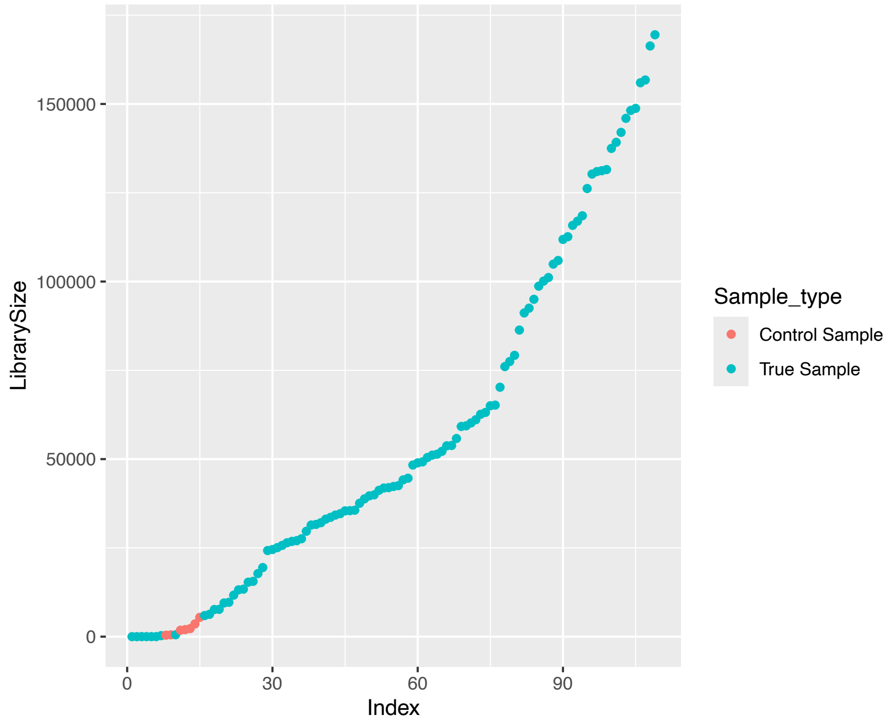

**B.**

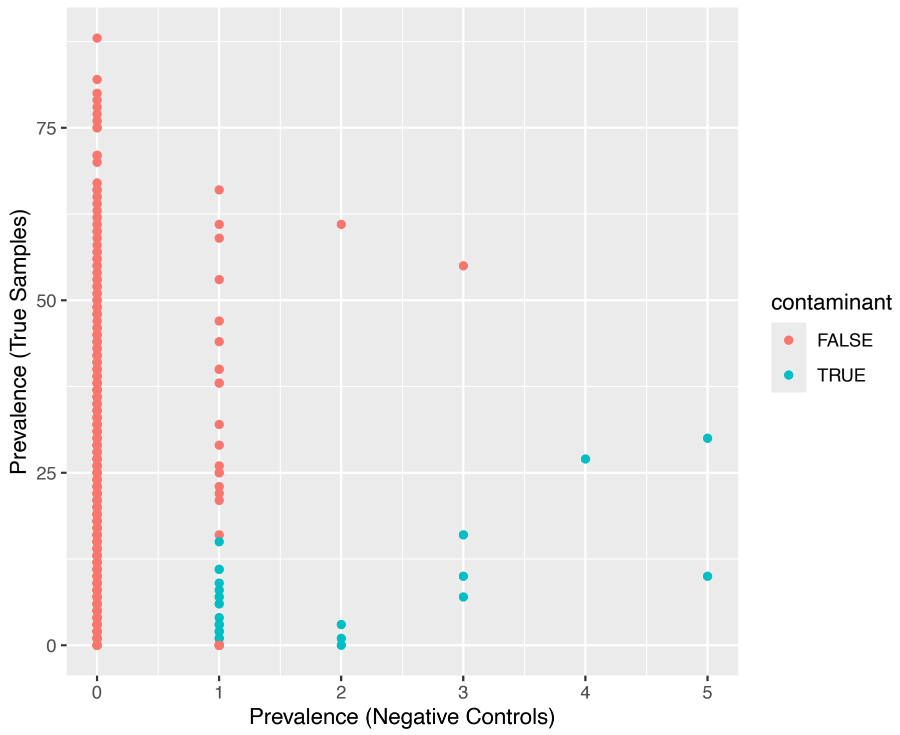

**C.**

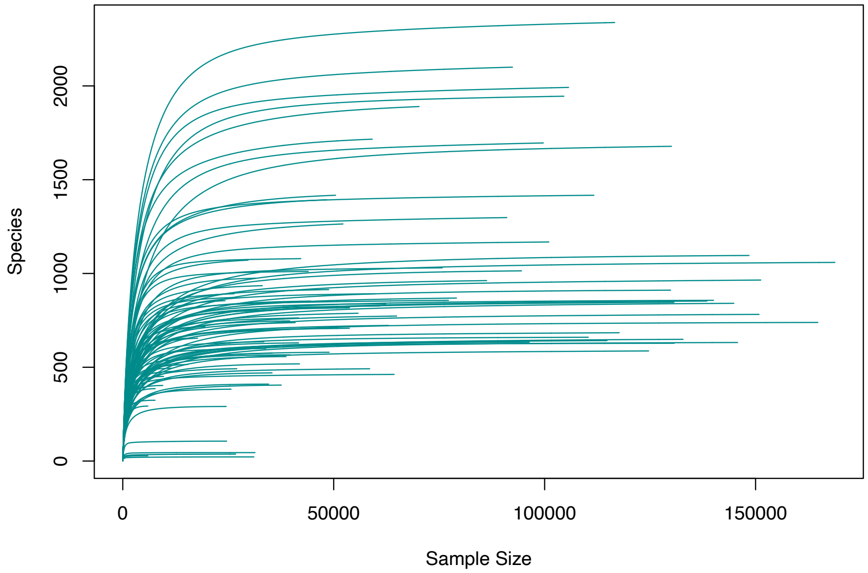

**Figure S1:** Decontamination diagnostics and sequencing depth. A. Library size of all samples, B. Prevalence of ASVs in negative controls and true samples, and C. Rarefaction curves for all samples.

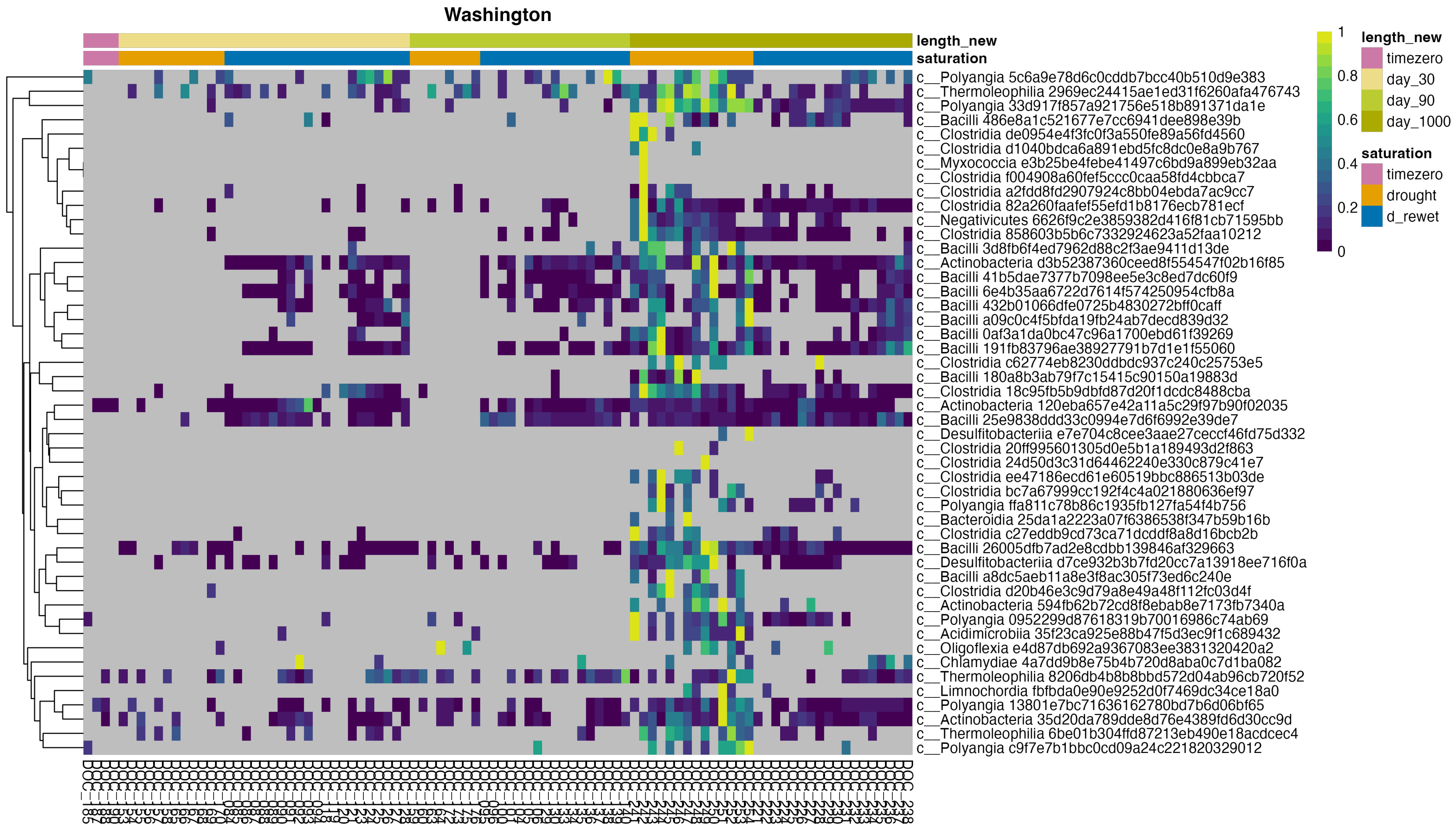

**Figure S2:** Forty-eight unique indicator taxa for Day 1000 drought.

**A.**

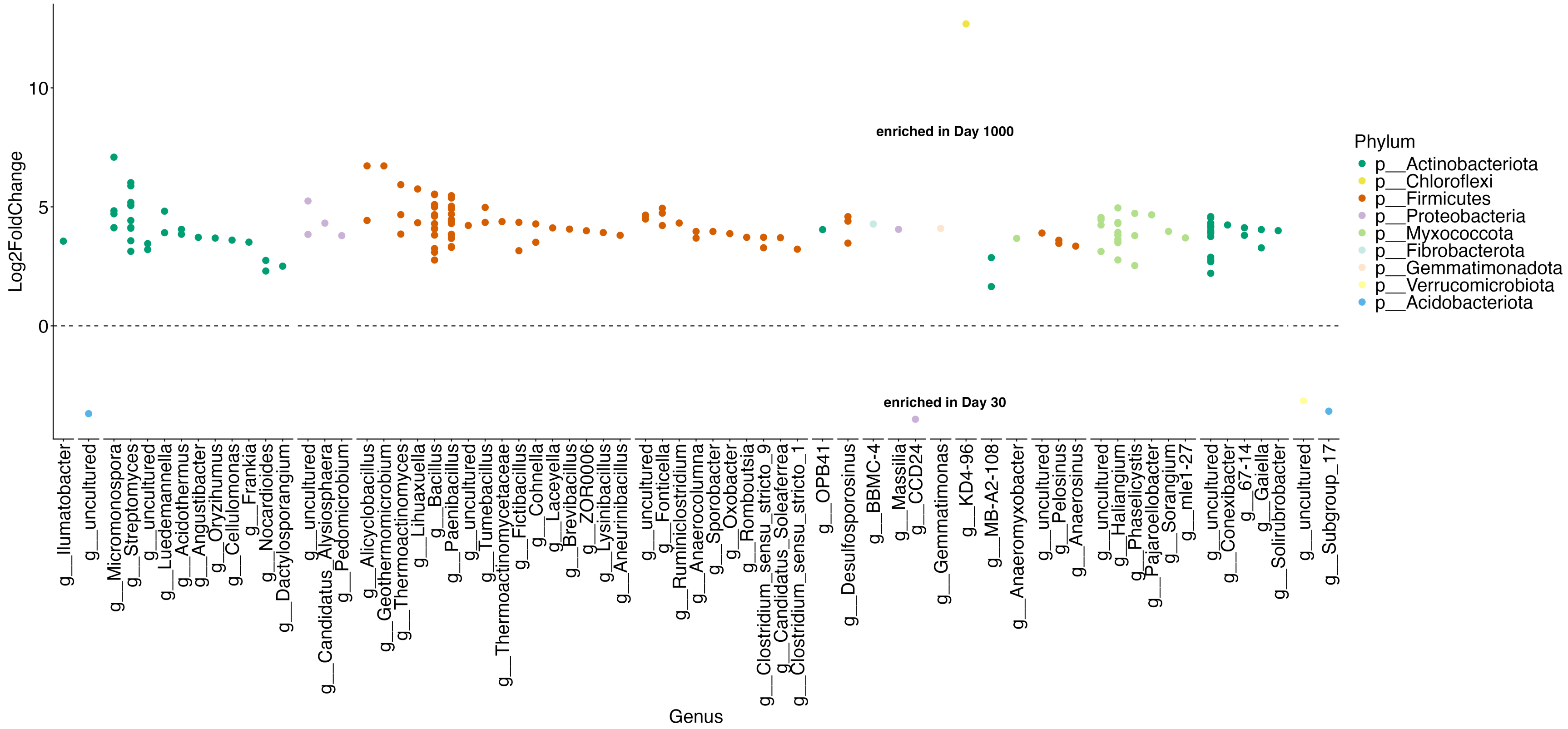
enriched in Day 1000
enriched in Day 30
Log2FoldChange
10
5
0
Genus
Phylum
p__Actinobacteriota
p__Chloroflexi
p__Firmicutes
p__Proteobacteria
p__Myxococcota
p__Fibrobacterota
p__Gemmatimonadota
p__Verrucomicrobiota
p__Acidobacteriota
g__Ilumatobacter
g__uncultured
g__Micromonospora
g__Streptomyces
g__uncultured
g__Luedemannella
g__Acidothermus
g__Angustibacter
g__Oryzihumus
g__Cellulomonas
g__Frankia
g__Nocardioides
g__Dactylosporangium
g__uncultured
g__Candidatus_Alysiosphaera
g__Pedomicrobium
g__Alicyclobacillus
g__Geothermomicrobium
g__Thermoactinomyces
g__Lihuaxuella
g__Bacillus
g__Paenibacillus
g__uncultured
g__Tumebacillus
g__Thermoactinomycetaceae
g__Fictibacillus
g__Cohnella
g__Laceyella
g__Brevibacillus
g__ZOR0006
g__Lysinibacillus
g__Aneurinibacillus
g__uncultured
g__Fonticella
g__Ruminiclostridium
g__Anaerocolumna
g__Sporobacter
g__Oxobacter
g__Romboutsia
g__Clostridium_sensu_stricto_9
g__Candidatus_Soleaferrea
g__Clostridium_sensu_stricto_1
g__OPB41
g__Desulfosporosinus
g__BBMC-4
g__Massilia
g__CCD24
g__Gemmatimonas
g__KD4-96
g__MB-A2-108
g__Anaeromyxobacter
g__uncultured
g__Pelosinus
g__Anaerosinus
g__uncultured
g__Haliangium
g__Phaselicystis
g__Pajaroellobacter
g__Sorangium
g__mle1-27
g__uncultured
g__Conexibacter
g__67-14
g__Gaiella
g__Solirubrobacter
g__uncultured
g__Subgroup_17

**B.**

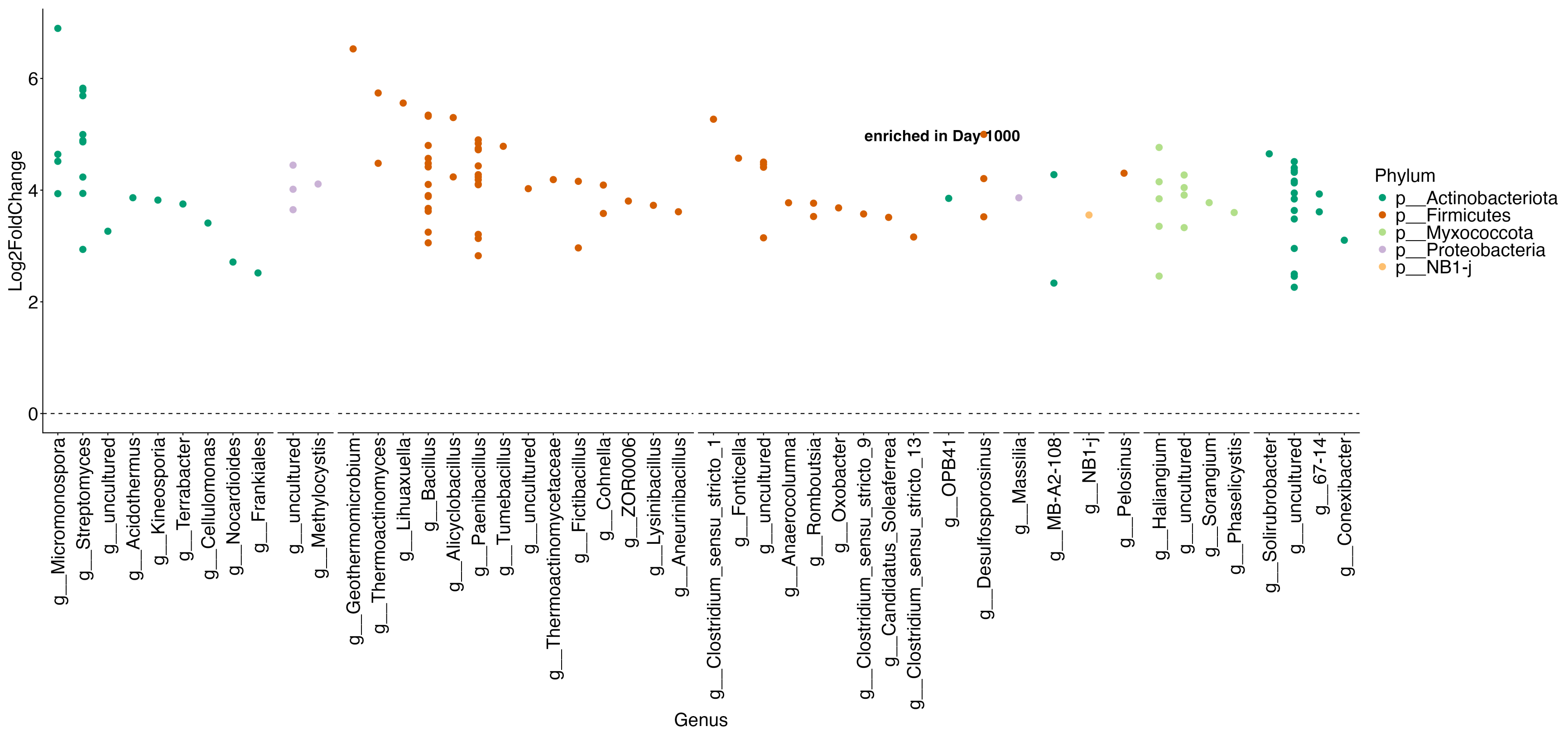

**Figure S3: Differential expression analysis on unrarefied data retaining all reads and samples to confirm enrichment of taxa at Day 1000 compared to A. Day 30 and B. Day 90.** Statistically significant enrichment of taxa belonging to Firmicutes and Actinobacteriota is observed at Day 1000 compared to Day 30 and Day 90 time points, confirming that there was no bias on the results due to rarefaction.

**A.**

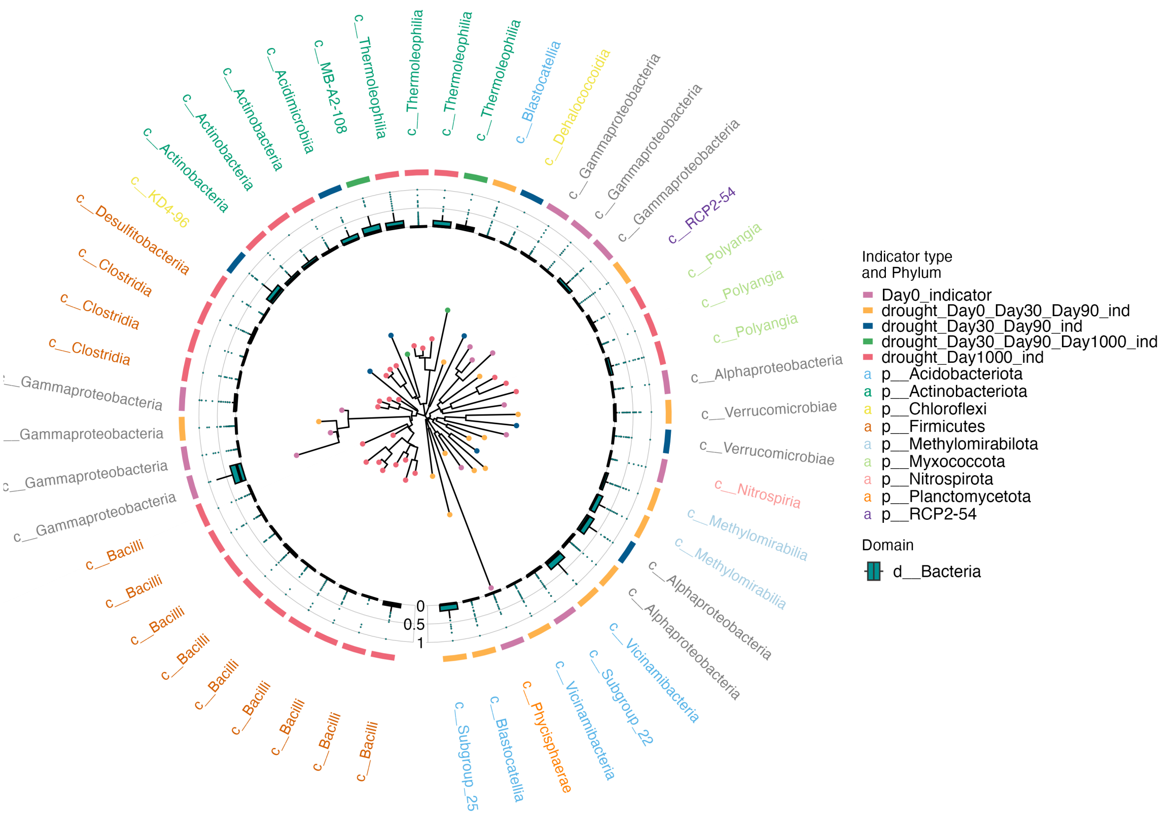
Indicator type
and Phylum
Day0_indicator
drought_Day0_Day30_Day90_ind
drought_Day30_Day90_ind
drought_Day30_Day90_Day1000_ind
drought_Day1000_ind
p__Acidobacteriota
p__Actinobacteriota
p__Chloroflexi
p__Firmicutes
p__Methylomirabilota
p__Myxococcota
p__Nitrospirota
p__Planctomycetota
p__RCP2-54
Domain
d__Bacteria
0
0.5
1
c__Thermoleophilia
c__Thermoleophilia
c__Thermoleophilia
c__Thermoleophilia
c__MB-A2-108
c__Acidimicrobiia
c__Actinobacteria
c__Actinobacteria
c__Actinobacteria
c__KD4-96
c__Desulfitobacteriia
c__Clostridia
c__Clostridia
c__Clostridia
c__Gammaproteobacteria
c__Gammaproteobacteria
c__Gammaproteobacteria
c__Gammaproteobacteria
c__Bacilli
c__Bacilli
c__Bacilli
c__Bacilli
c__Bacilli
c__Bacilli
c__Bacilli
c__Bacilli
c__Bacilli
c__Subgroup_25
c__Blastocatellia
c__Phycisphaerae
c__Vicinamibacteria
c__Subgroup_22
c__Vicinamibacteria
c__Alphaproteobacteria
c__Alphaproteobacteria
c__Methylomirabilia
c__Methylomirabilia
c__Nitrospiria
c__Verrucomicrobiae
c__Verrucomicrobiae
c__Alphaproteobacteria
c__Polyangia
c__Polyangia
c__Polyangia
c__RCP2-54
c__Gammaproteobacteria
c__Gammaproteobacteria
c__Gammaproteobacteria
c__Dehalococcoidia
c__Blastocatellia

**B.**

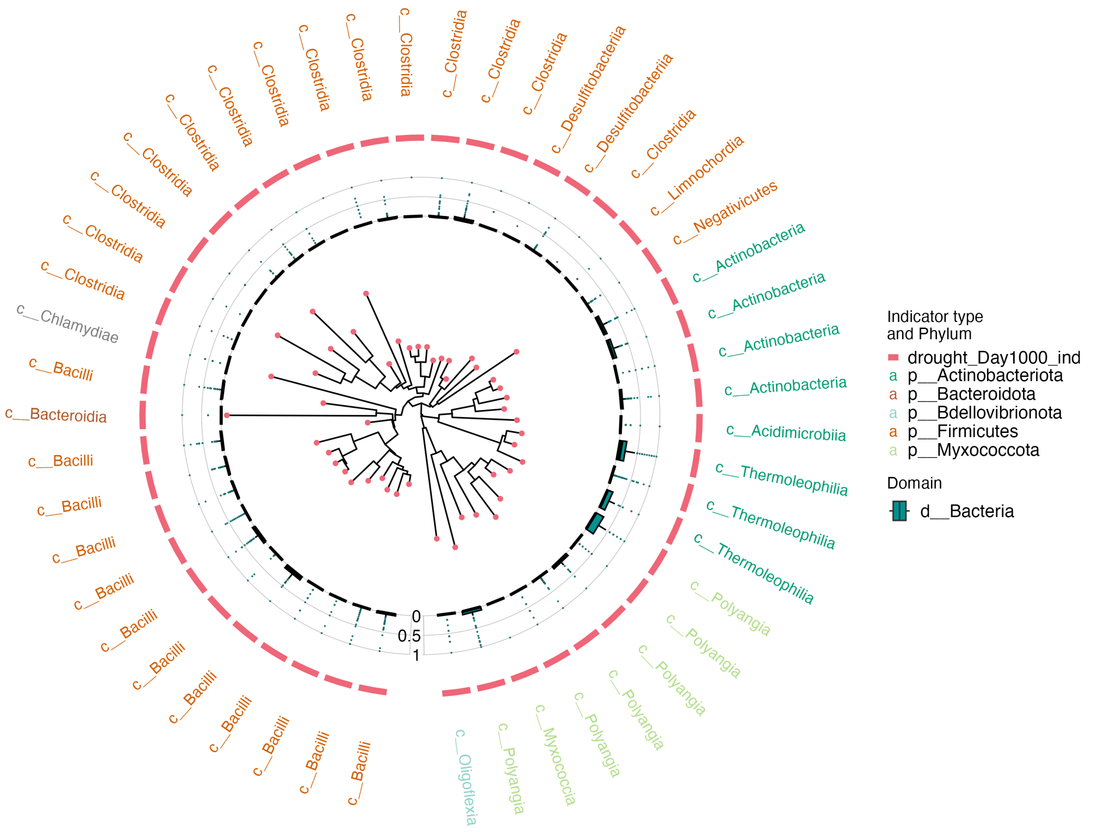

**Figure S4**: **Drought responses are conserved at higher taxonomic levels across short-term and long-term drought. A.** Phylogenetic tree or dendrogram showing the top 50 indicator taxa (corresponding to heatmap Fig 3B) for all drought timepoints (Day 0, Day 30, Day 90, Day 1000). **B.** Dendrogram for unique drought responsive taxa at Day 1000 (48 total).

**A.**

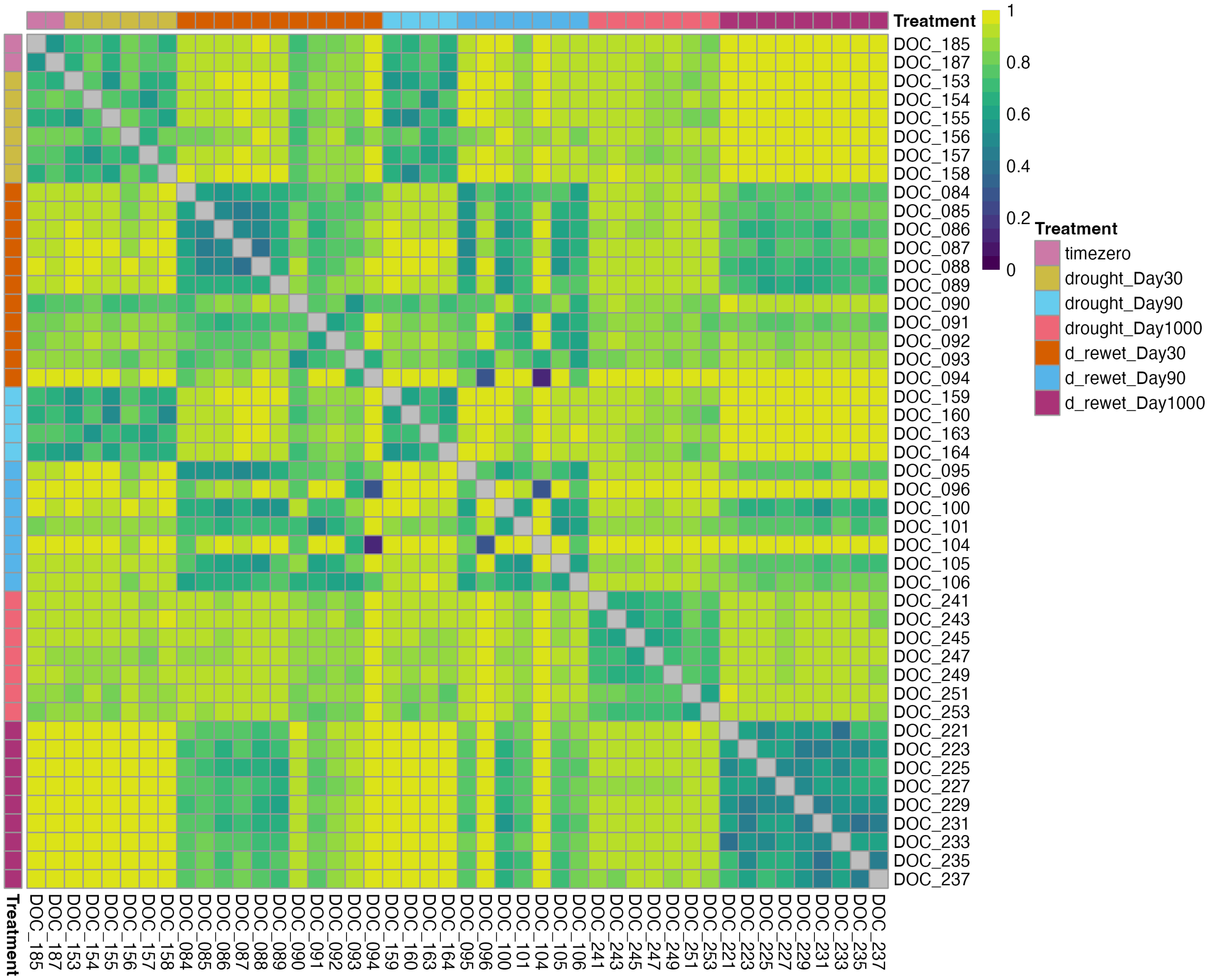
Treatment
1
0.8
0.6
0.4
0.2
0
Treatment
timezero
drought_Day30
drought_Day90
drought_Day1000
d_rewet_Day30
d_rewet_Day90
d_rewet_Day1000
DOC_185
DOC_187
DOC_153
DOC_154
DOC_155
DOC_156
DOC_157
DOC_158
DOC_084
DOC_085
DOC_086
DOC_087
DOC_088
DOC_089
DOC_090
DOC_091
DOC_092
DOC_093
DOC_094
DOC_159
DOC_160
DOC_163
DOC_164
DOC_095
DOC_096
DOC_100
DOC_101
DOC_104
DOC_105
DOC_106
DOC_241
DOC_243
DOC_245
DOC_247
DOC_249
DOC_251
DOC_253
DOC_221
DOC_223
DOC_225
DOC_227
DOC_229
DOC_231
DOC_233
DOC_235
DOC_237

**B.**

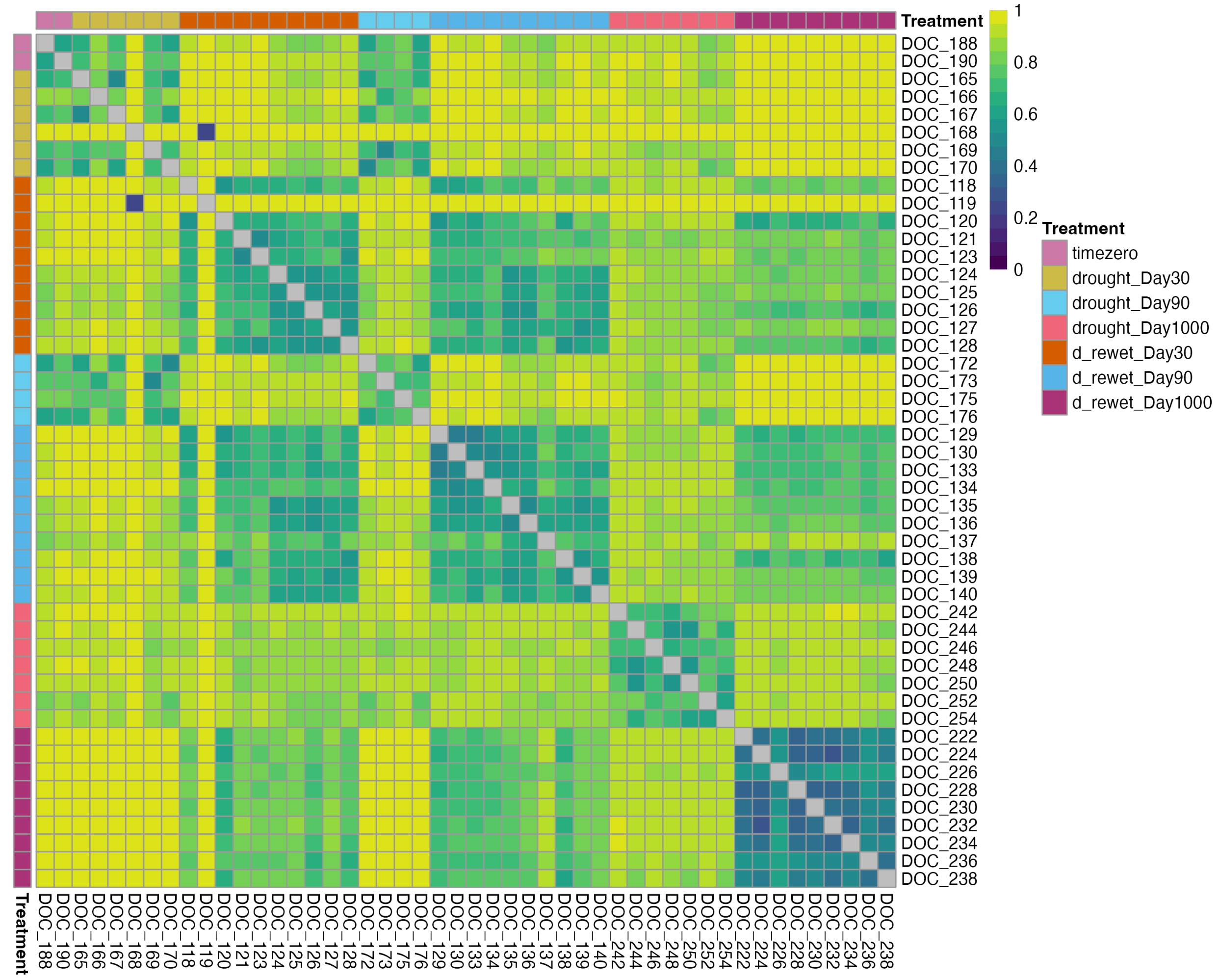

**Figure S5: Bray-Curtis distances between drought and drought-rewet samples at 30d, 90d and 1000d time points.** Color gradient denotes Bray-Curtis distance dissimilarity index ranging from 0 (100% similarity - dark blue color in color gradient) to 1 (100% dissimilarity - bright yellow color in color gradient). A. 0-5 cm depth, B. 5cm-end depth.

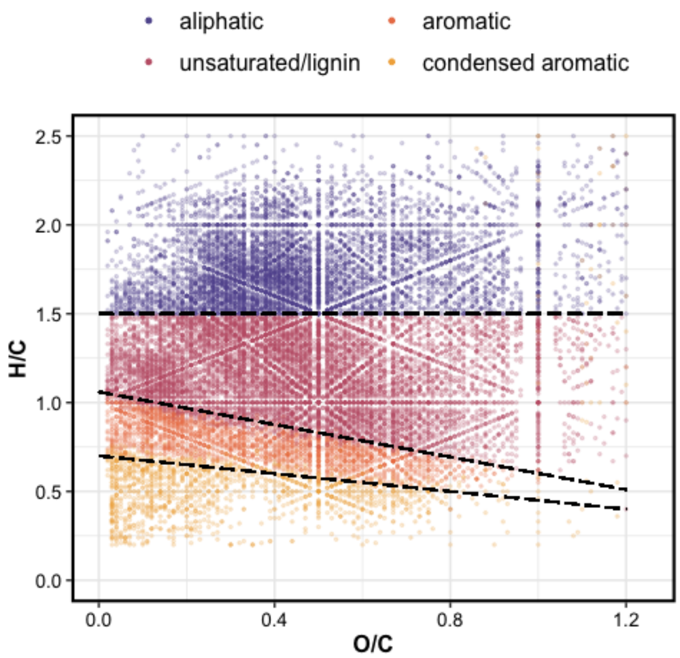

**Figure S6:** Van Krevelen diagram, with molecular classification of the peaks identified via FT-ICR-MS. Each point represents a unique peak, plotted against its molecular H/C and O/C ratios.

**A.**

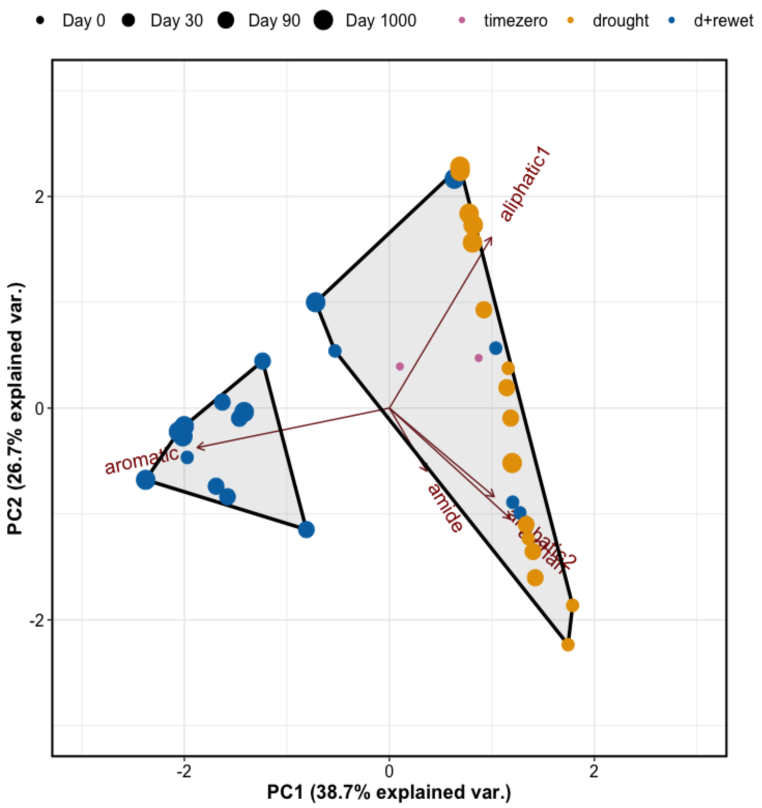
Day 0
Day 30
Day 90
Day 1000
timezero
drought
d+rewet
aliphatic1
aromatic
amide
2
0
-2
-2
0
2
PC1 (38.7% explained var.)
PC2 (26.7% explained var.)

**B.**

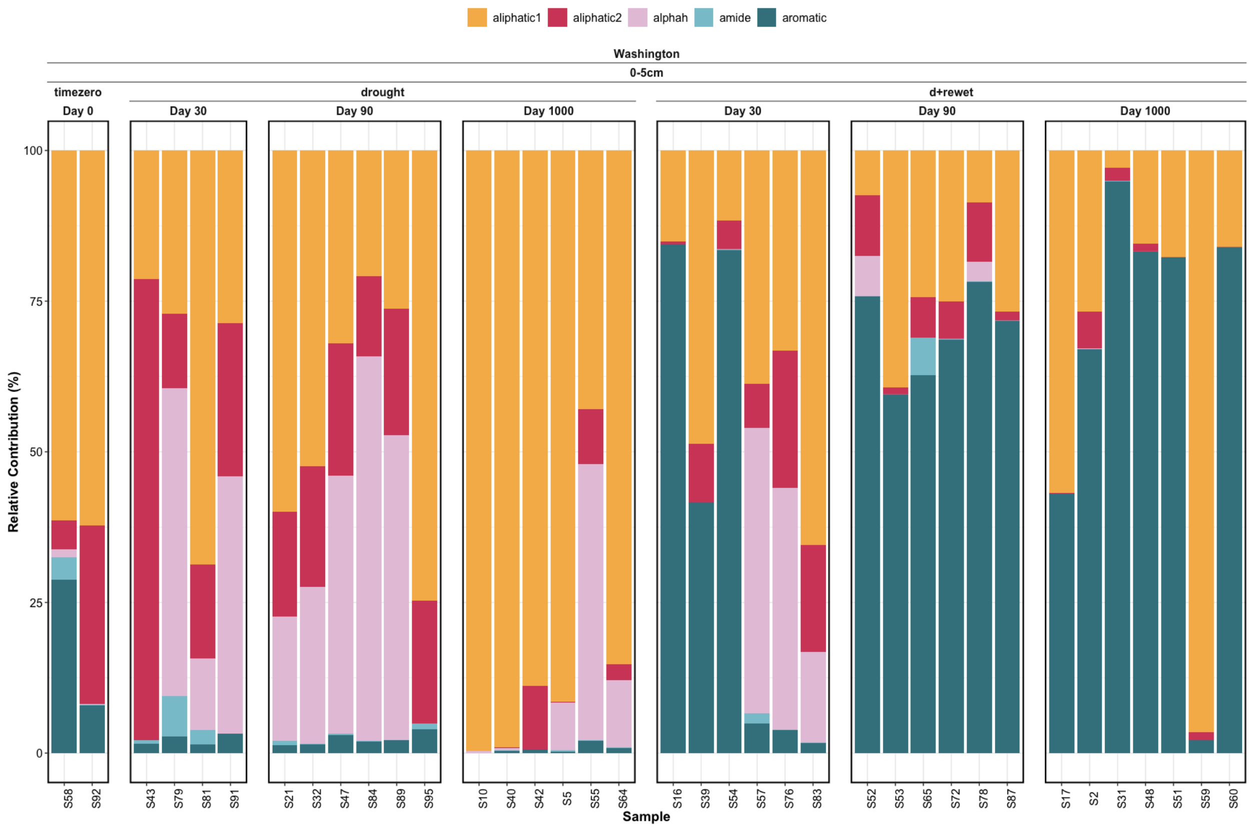
aliphatic1
aliphatic2
alphah
amide
aromatic
Washington
0-5cm
timezero
drought
d+rewet
Day 0
Day 30
Day 90
Day 1000
Relative Contribution (%)
Sample
S58 S92
S43 S79 S81 S91
S21 S32 S47 S84 S89 S95
S10 S40 S42 S5 S55 S64
S16 S39 S54 S57 S76 S83
S52 S53 S65 S72 S78 S87
S17 S2 S31 S48 S51 S59 S60

**C.**

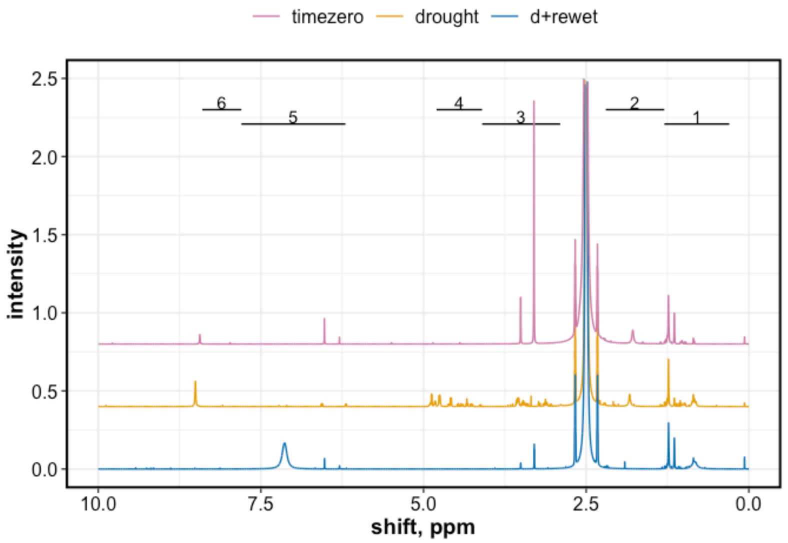
timezero
drought
d+rewet
intensity
shift, ppm
1
2
3
4
5
6

**Figure S7**: **NMR PCA, barplots, and example spectra. A** PCA biplot of NMR data. The data are grouped into two clusters (identified via hierarchical clustering) – one cluster contains all timezero and drought samples, whereas the other cluster contains all d+rewet samples. **B**. Relative abundance of NMR-resolved functional groups. Drought samples showed increased alpha-H groups (protein), whereas d+rewet samples showed increased aromatic groups, compared to the time-zero samples. **C.** Example NMR spectra (one from each treatment type). Compared to timezero, the drought samples showed unique peaks in the alpha-H/protein region (#4), and the d+rewet samples showed greater abundance of peaks in the aromatic region (#5).

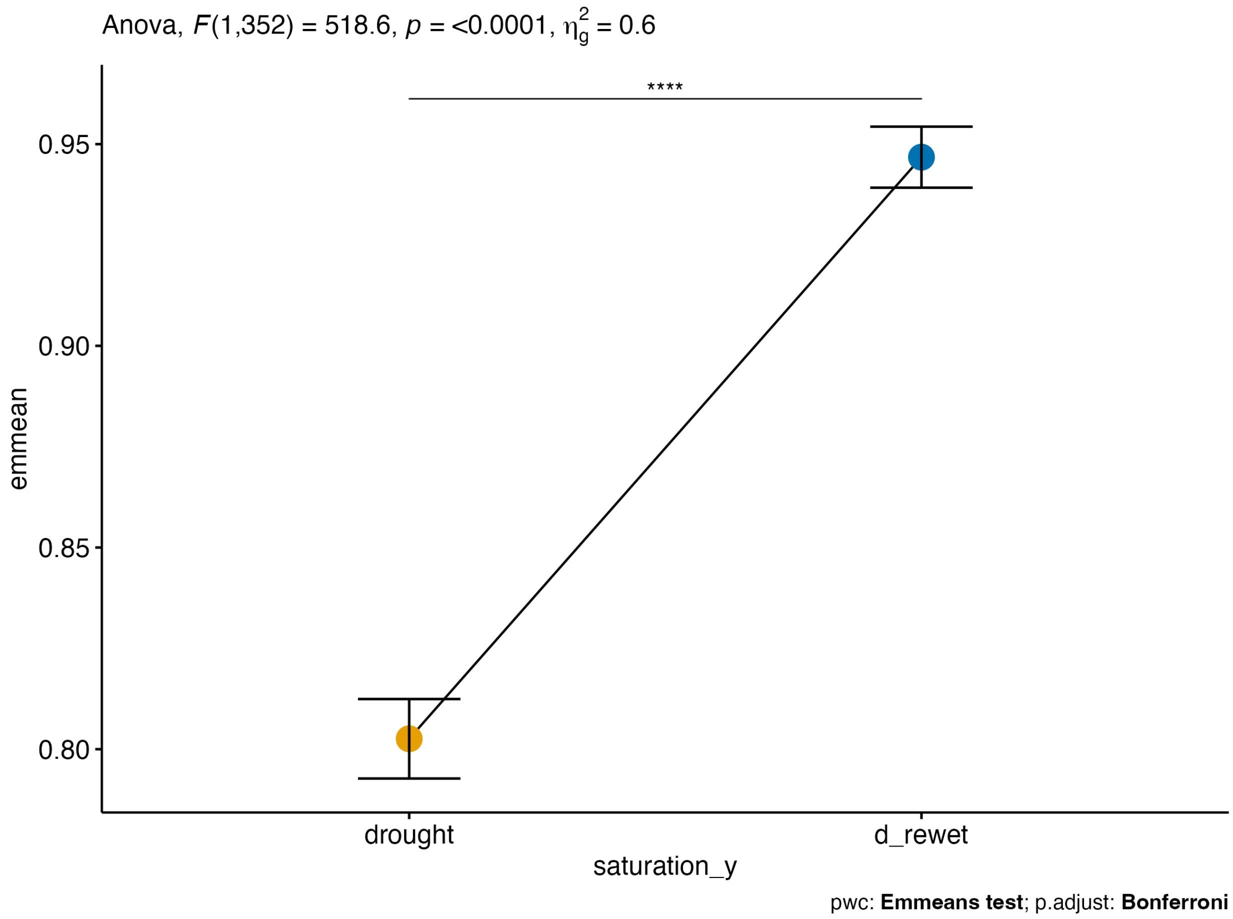

**Figure S8**: Analysis of covariates. Difference between the linear trends in Bray Curtis similarity to pre-drought (Day 0) for drought and d+rewet samples (Fig 7, main manuscript), accounting for covariate of time
